# The extracellular gate shapes the energy profile of an ABC exporter

**DOI:** 10.1101/421073

**Authors:** Cedric A.J. Hutter, M. Hadi Timachi, Lea M. Hürlimann, Iwan Zimmermann, Pascal Egloff, Hendrik Göddeke, Svetlana Kucher, Saša Štefanic, Mikko Karttunen, Lars V. Schäfer, Enrica Bordignon, Markus A. Seeger

**Affiliations:** Institute of Medical Microbiology, University of Zurich, Gloriastr. 28/30, 8006 Zurich, Switzerland; Faculty of Chemistry and Biochemistry, Ruhr University Bochum, 44801 Bochum, Germany; Institute of Parasitology, University of Zurich, Winterthurerstrasse 266a, 8057 Zurich, Switzerland; Theoretical Chemistry, Faculty of Chemistry and Biochemistry, Ruhr University Bochum, 44801 Bochum, Germany; Department of Chemistry and Department of Applied Mathematics, The University of Western Ontario, London, Ontario N6A 3K7, Canada

## Abstract

ABC exporters harness the energy of ATP to pump substrates across membranes. Extracellular gate opening and closure are key steps of the transport cycle, but the underlying mechanism is poorly understood. Here, we generated a synthetic single domain antibody (sybody) that recognizes the heterodimeric ABC exporter TM287/288 exclusively in the presence of ATP, which was essential to solve a 3.2 Å crystal structure of the outward-facing transporter. The sybody binds to an extracellular wing and strongly inhibits ATPase activity by shifting the transporter’s conformational equilibrium towards the outward-facing state, as shown by double electron-electron resonance (DEER). Mutations that facilitate extracellular gate opening resulted in a comparable equilibrium shift and strongly reduced ATPase activity and drug transport. Using the sybody as conformational probe, we demonstrate that efficient extracellular gate closure is required to dissociate the NBD dimer after ATP hydrolysis to reset the transporter back to its inward-facing state.

## INTRODUCTION

ABC exporters are versatile membrane proteins found in all phyla of life. Type I exporters are the best studied class of ABC exporters and minimally consist of two transmembrane domains (TMDs) each comprising six transmembrane helices and two nucleotide binding domains (NBDs) that are universally conserved among all ABC transporters. The NBDs undergo large conformational changes in response to ATP binding and hydrolysis, which are transmitted to the TMDs via coupling helices to assume inward-facing (IF), outward-facing (OF) and outward-occluded (Occ) conformations^1^. Alternating access at the TMDs in conjugation with affinity changes towards the transported substrates enable uphill transport across the lipid bilayer^2^. Fully closed NBDs are stabilized by two ATP molecules bound at the dimer interface and coincide with TMDs adopting an OF or Occ state^3, 4^. The transition to the IF state requires the NBDs to separate at least to some degree, a process that is initiated by ATP hydrolysis^5^.

Many ABC exporters including the entire human ABCC family exhibit asymmetric ATP binding sites, namely a degenerate site that can bind but not hydrolyze ATP and a consensus site that is hydrolysis-competent^6^. Heterodimeric TM287/288 of the thermophilic bacterium *Thermotoga maritima* was the first structurally analyzed example of an ABC exporter with a degenerate site^7, 8^. Two closely related IF structures of TM287/288 were solved by X-ray crystallography either containing one AMP-PNP molecule bound to the degenerate site or no nucleotide. In contrast to most other IF structures of ABC exporters, the opened NBDs of TM287/288 are only partially separated due to contacts mediated by the degenerate site D-loop, whereas the consensus site D-loop was found to allosterically couple ATP binding at the degenerate site to ATP hydrolysis at the consensus site^8^. The consensus site features distortions in the Walker B motif, which prevents nucleotide binding in the IF transporter^7^. DEER studies have revealed that TM287/288 exhibits dynamic IF/OF equilibria in the presence of nucleotides and that nucleotide trapping at the consensus site is required to strongly populate the OF state, whereas in the presence of AMP-PNP the transporter predominantly adopts its IF state^9^.

Broad distance distributions were found by DEER in the extracellular gate of TM287/288, hinting at conformational flexibility in this external region^9^. Similar observations were reported for ABCB1^10^. Unbiased MD simulations of TM287/288 uncovered spontaneous conformational transitions from the IF state via an Occ intermediate to the OF state^11^. Many simulations remained trapped in the Occ state, suggesting that extracellular gate opening represents a major energetic barrier in the conformational cycle. Interestingly, the degree of extracellular gate opening varies greatly among different type I ABC exporters solved in OF states, whereas the gate remains closed in the Occ state^3, 4, 12^. Hence, events occurring at the extracellular gate likely play a key role in substrate transport and must be allosterically coupled to the catalytic cycle of the NBDs. Nevertheless, the underlying molecular mechanism is unknown.

In this work, we generated single domain antibodies that exclusively bind to outward-facing TM287/288 and thereby inhibit the transport cycle. The binders were instrumental to solve a crystal structure of the transporter in its OF state and were used to probe molecular events at the extracellular gate and their allosteric coupling with the NBDs.

## RESULTS

### Conformational trapping of TM287/288

Having solved two closely related IF structures of TM287/288, our aim was to obtain an atomic structure of this heterodimeric ABC exporter in its OF state. DEER analyses revealed that TM287/288 carrying the TM288^E^517^Q^ mutation in the Walker B motif of the consensus site (EtoQ mutation) was almost completely trapped in the OF state in the presence of ATP-Mg and ATPγS-Mg^9^. To further decrease the residual ATPase activity of the EtoQ mutant (turnover of 0.02 min^-1^) by a factor of 6.5, we instead introduced the EtoA mutation. In addition, we generated single domain antibodies (nanobodies) that exclusively recognize the OF state of TM287/288. To this end, alpacas were immunized with outward-facing TM287/288 containing a cross-linked tetrahelix bundle motif^13^ (see Materials and Methods). This approach yielded nanobody Nb_TM#1 binding exclusively to TM287/288 in the presence (but not in the absence) of ATP, as shown by surface plasmon resonance (SPR) (Fig. 1d). However, crystals obtained with Nb_TM#1 did not diffract well enough to build a reliable model. Therefore, we selected synthetic nanobodies (sybodies) against TM287/288(EtoA) in the presence of ATP-Mg completely *in vitro*^14^. *Thereby, more than ten OF-specific sybodies were generated and sybody Sb_TM#35 was successfully used to solve the OF structure of TM287/288(EtoA) in the presence of ATPγS-Mg at 3.2 Å resolution (Fig. 1a, Table S1).*

**Figure 1.**
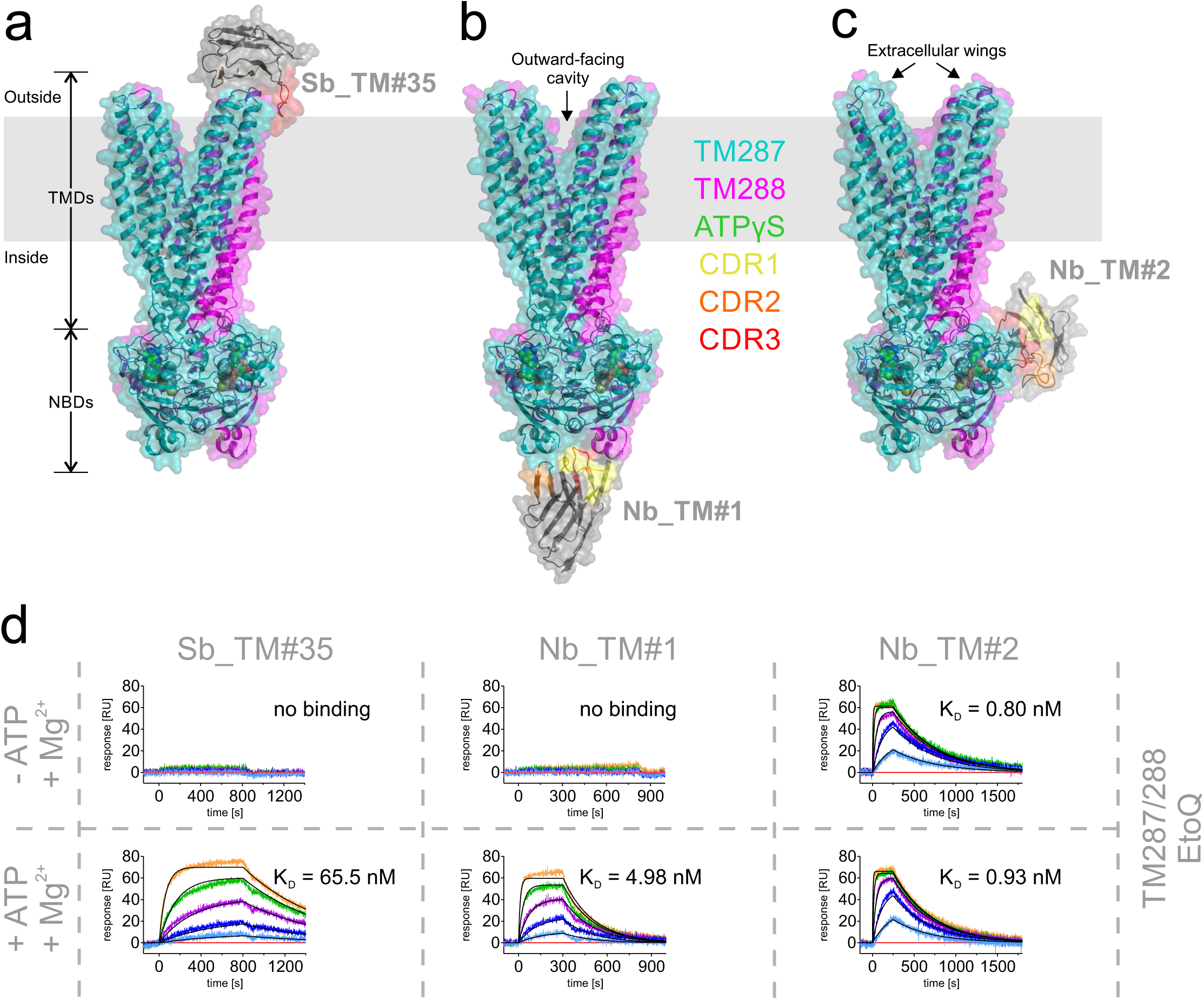
Three outward-facing structures of TM287/288 solved in complex with single domain antibodies. The transporters are viewed along the membrane plane (indicated as grey rectangle). (**a**) 3.2 Å structure of TM287/288(EtoA) solved in complex with ATPγS-Mg and state-specific sybody Sb_TM#35. (**b**) 3.5 Å structure of TM287/288(2xDtoA/EtoA) in complex with ATPγS-Mg and state-specific nanobody Nb_TM#1. (**c**) 4.2 Å crystal structure of TM287/288(2xDtoA/EtoA) in complex with ATPγS-Mg and state-unspecific nanobody Nb_TM#2. (**d**) SPR analyses in the presence (upper panel) and absence (lower panel) of ATP using immobilized TM287/288(EtoQ) as ligand and Sb_TM#35, Nb_TM#1 and Nb_TM#2 as analytes. Injected concentrations of Sb_TM#35: 0, 9, 27, 81, 243, 729 nM; Nb_TM#1: 0, 1, 3, 9, 27, 81 nM; Nb_TM#2: 0, 0.9, 2.7, 8.1, 24.3, 72.9 nM. Kinetic analysis is shown in Table S2.

### Structure of OF TM287/288 with a sybody bound to an extracellular wing

Sybody Sb_TM#35 binds on top of an extracellular wing of TM287/288 (Fig. 1a) and was crucially involved in establishing crystal contacts (Fig. S1). Binding is mediated by aromatic residues of all three complementary determining regions (CDRs) of the sybody, which are wedged between transmembrane helices (TMs) 1 and 2 of TM287 and TMs 5’ and 6’ of TM288 (Fig. 2a). Since Sb_TM#35 only binds in the presence of ATP (Fig. 1d), we hypothesized that it interferes with the catalytic cycle of the transporter. Indeed, the sybody inhibited the ATPase activity of TM287/288 in detergent (IC_50_ of 66.1 nM, Fig. 2b) as well as reconstituted in nanodiscs (Fig. S2b). Hence, the sybody addresses an epitope that is accessible by biopharmaceuticals in the cellular context.

**Figure 2.**
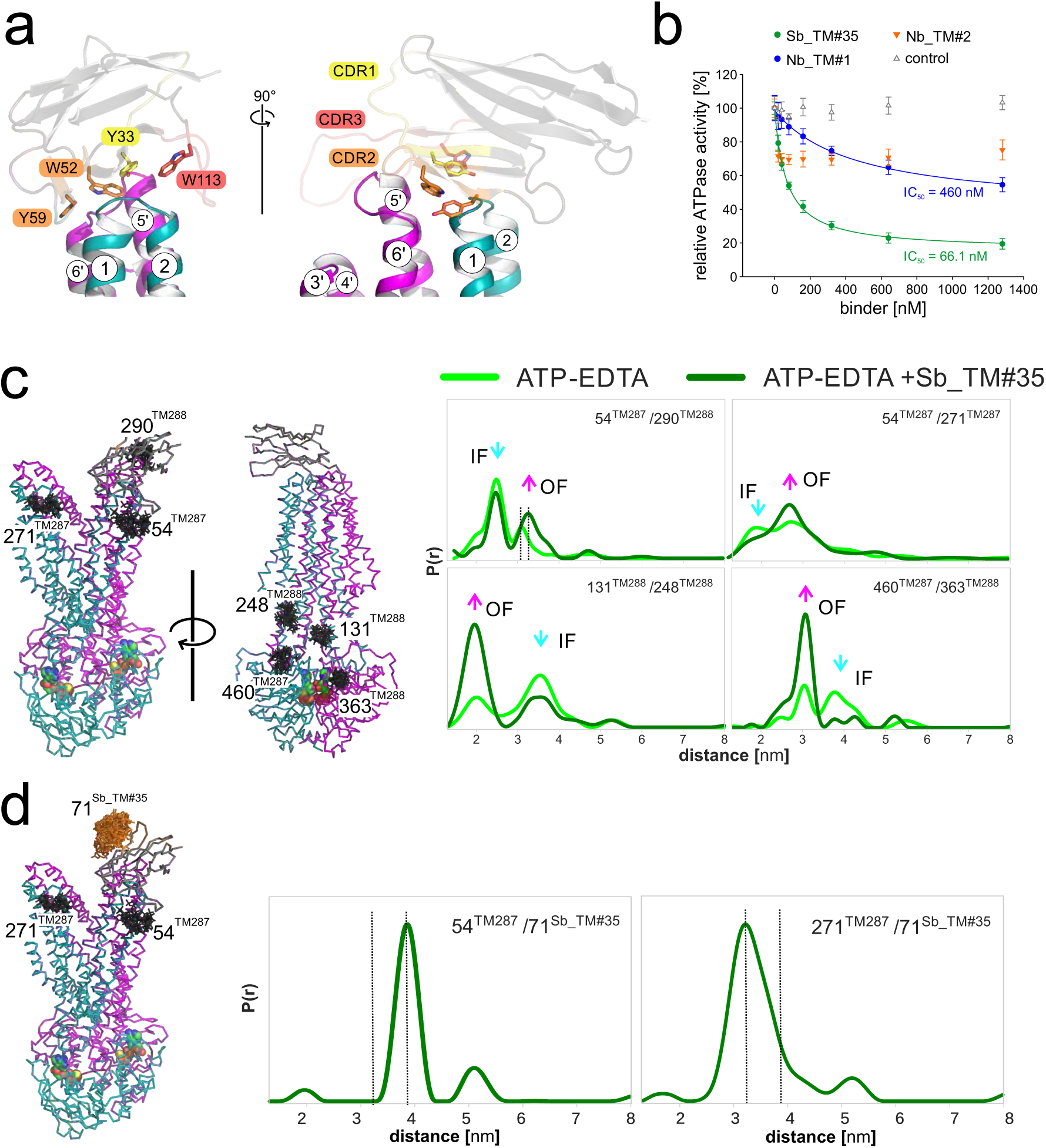
The sybody traps TM287/288 in its OF state. (**a**) Sybody Sb_TM#35 is shown as cartoon in grey with the CDR1, 2 and 3 highlighted in yellow, orange and red, respectively. Four aromatic residues (Y33, W52, Y59 and W113) that wedge between TMs 1 and 2 of TM287 (teal) and TMs 5’ and 6’ of TM288 (magenta) are highlighted as sticks. (**b**) Inhibition of TM287/288’s ATP hydrolysis by Sb_TM#35, Nb_TM#1 and Nb_TM#2. A non-randomized sybody served as control. The data were fitted with a hyperbolic decay function to determine IC_50_ values as well as residual activities. The error bars are standard deviations of technical triplicates. (**c** and **d**) DEER analysis of spin-label pairs introduced to probe the extracellular and intracellular TMDs and the NBDs (**c**) as well as sybody binding to the transporter (**d**). DEER traces were recorded in the presence of ATP-EDTA with or without adding unlabeled Sb_TM#35 (**c**) or with spin-labeled Sb_TM#35 (**d**) and the graphs show experimental distance distributions.

### Two nanobodies addressing epitopes on the NBDs

Using the high resolution structure of outward-facing TM287/288 for molecular replacement and as template for model building, we solved two additional low resolution structures (3.5 - 4.2 Å) of the OF transporter determined in complex with alpaca nanobodies Nb_TM#1 and Nb_TM#2 (Fig. 1b and c). Nb_TM#1 binds to the bottom of the closed NBD dimer and occupies an epitope that is shared between NBD287 and NBD288, which explains why it specifically recognizes the OF state (Fig. 1d). Akin to Sb_TM#35, state-specific Nb_TM#1 was found to inhibit the transporter’s ATPase activity (Fig. 2b). Nb_TM#2 binds side-ways to NBD288 and exhibits picomolar affinity for the transporter regardless whether ATP is present or not (Fig. 1d, Table S2). Nevertheless, this nanobody partially inhibits ATPase activity by around 30% already at very low concentrations, which prevented us from determining an IC_50_ value (Fig. 2b). A measurement artifact can be excluded, because an unrelated control sybody did not affect the transporter’s ATPase activity (Fig. 2b).

### Transition from IF to OF state renders TM287/288 more symmetric

The OF structure of TM287/288 features fully dimerized NBDs that sandwich two ATP molecules at the degenerate and the consensus site (Fig. 3a). In contrast to the NBDs of inward-facing TM287/288, which exhibited pronounced asymmetries between degenerate and consensus site mainly with regard to the D-loops^8^, the closed NBD dimer of the OF transporter is more symmetric (Fig. 3a, Fig. S3). Further, the distortions found at the catalytic dyad of the consensus site of the IF structure (E517^TM288^ and H548^TM288^) relax during the transition to the OF state and the two key residues adopt a hydrolysis-competent arrangement (Fig. 3b). Interestingly, two tunnels that would allow for release of the cleaved γ-phosphate are present at the consensus site (Fig. 3c). The TMDs consisting of two wings each encompassing six transmembrane helices donated from both protomers is widely opened towards the outside (Fig. S4). With a RMSD of 1.73 Å, the structure of TM287/288 most closely resembles the structure of Sav1866 and is similar to the OF state of TM287/288 predicted by MD simulations (Fig. S5)^11^. Also the degree of NBD closure and extracellular gate opening is highly similar between TM287/288 and Sav1866 (Fig. S6a). The RMSD between TM287 and TM288 decreases from 2.55 Å to 1.98 Å as the transporter is converted from the IF to the OF conformation, indicating that outward-facing TM287/288 is more symmetric (Fig. S6b). Whereas a similar degree of symmetry was observed between the half-transporters of outward-facing ABCB1 (PDB: 6C0V, RMSD of 2.07 Å), the equivalent superimpositions exhibit substantial asymmetries in the OF structure of MRP1 (PDB: 6BHU, RMSD of 4.54 Å), mostly owing to asymmetries in the TMDs (Fig. S6b). Extracellular gate opening is less pronounced in MRP1 and even less so in ABCB1, and the gate remains almost completely closed in the outward-occluded (Occ) structure of McjD^4^ (Fig. S6b). Hence, structures of OF and Occ ABC exporters show their largest structural variation in the extracellular gates.

**Figure 3.**
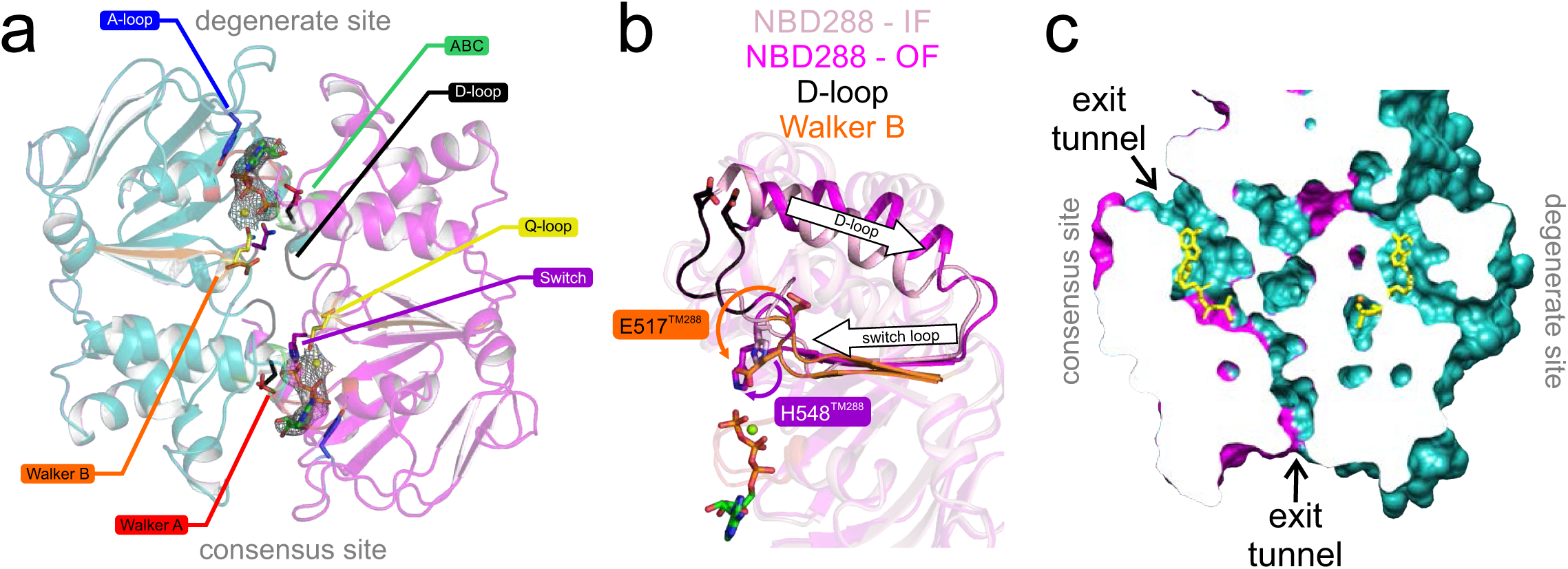
Structural analysis of the closed NBD dimer. (**a**) The fully closed NBD dimer (NBD287 in teal and NBD288 in magenta) sandwiches two ATPγS-Mg molecules (shown as sticks with corresponding electron density) between Walker A motif (red) and the opposite ABC signature motif (green) at the degenerate and the consensus site in a highly symmetric manner. Residues involved in ATP binding and hydrolysis are shown as sticks. (**b**) Superimposition of the consensus ATP binding site of the previously solved IF structure (PDB: 4Q4H, light pink) and the OF structure (magenta). Distortions of the catalytic dyad (E517^TM288^ and H548^TM288^) are relaxed during NBD closure to adopt a hydrolysis-competent arrangement. (**c**) Slice-cut through the two nucleotide binding sites reveals two possible P_i_ exit tunnels at the consensus site which are not present at the degenerate site. ATPγS (partially clipped) is shown as yellow sticks.

### The sybody acts as molecular clamp

Interestingly, we did not find steric clashes which would prevent Sb_TM#35 from binding to the IF transporter. Hence, based on structural information alone we could not explain why the sybody inhibits ATPase activity. Therefore, we used DEER spectroscopy to unravel the sybody’s impact on the conformational cycle.

The sybody was found to shift the transporter’s equilibrium towards the OF state, as measured in the presence of ATP-EDTA (arrows in Fig. 2c). Pronounced effects were observed in the extracellular region (54^TM287^/290^TM288^ and 54^TM287^/271^TM287^), but also when probing distances at the intracellular region of the TMDs (131^TM288^/248^TM288^) and at the NBDs (460^TM287^/363^TM288^) (Fig. 2c and Fig. S7). Further, we observed a distance increase between two spin labels positioned in the wing underneath the sybody (54^TM287^/290^TM288^) as a result of sybody binding (dotted vertical lines in Fig. 2c). As expected from the lack of sybody binding to the IF state, we observed negligible effects on the interspin distances when TM287/288 was incubated with the sybody in the absence of nucleotides (apo state) (Fig. S7).

To investigate the positioning of the bound sybody relative to the opposite wing, we then focused on the distance between the sybody labeled at position 71 and spin-labels introduced either at 54^TM287^ (the sybody-binding wing) or 271^TM287^ (opposite wing) of the transporter (Fig. 2d and Fig. S8). The main distance peak corresponding to dipolar coupling between 71^Sb_TM#35^ and 54^TM287^ was very sharp and centered at 3.8 nm, while it was somewhat broader and centered at 3.2 nm between 71^Sb_TM#35^ and 271^TM287^ placed on the opposite wing. Both distances were visible only in the presence of ATP, and were in close agreement with the simulations based on the OF structure (Fig. S8). Both traces also contained a distance peak at around 5.2 nm corresponding to a residual fraction of sybody dimers. This peak was the only distance observed in control experiments examining the spin-labeled sybody alone or in the presence of the apo transporter (data not shown). In conclusion, the sybody acts as a molecular clamp that keeps the extracellular gate open.

### Conserved aspartates seal the extracellular gate

Having shown that the sybody traps the transporter in a fully opened state, we reasoned that mutations facilitating extracellular gate opening would have a similar impact on the transporter’s energy landscape. In inward-facing TM287/288, D41^TM287^ and D65^TM288^ placed in TM1 of the respective half-transporter establish hydrogen bonds with backbone amides of the opposite wing (Fig. 4a). Of note, these aspartates are conserved in bacterial ABC exporters (Fig. 4b). When the aspartates were mutated into alanines, the ATPase activity of TM287/288 decreased around three-fold for the single mutants and around 10-fold for the double mutant (henceforth called 2xDtoA mutant) (Fig. 4c).

**Figure 4.**
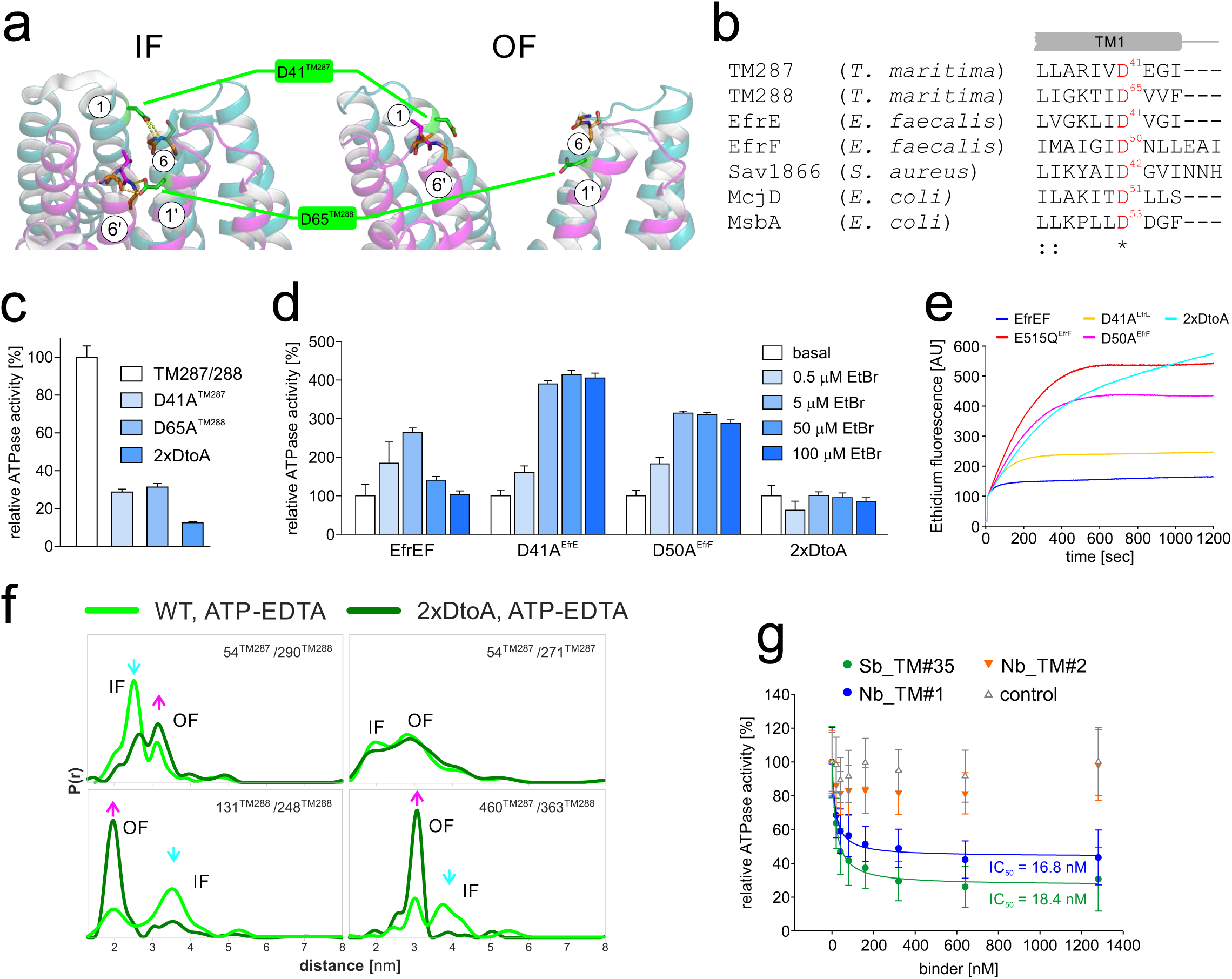
The extracellular gate is sealed by two conserved aspartates. (**a**) Structure of TM287/288’s extracellular gate in the IF (left, PDB: 4Q4H) and OF (right) state shown as cartoon. D41^TM287^and D65^TM288^ are shown as sticks and establish hydrogen bonds (dashed yellow lines) with the peptide backbone (shown as sticks) of neighboring helices that are broken during IF-OF transition. (**b**) Alignment of bacterial ABC exporters in the region containing the conserved extracellular gate aspartates. (**c**) ATPase activities of single mutants D41A^TM287^ and D65A^TM288^ and the corresponding double mutant (2xDtoA) relative to wildtype TM287/288 determined in detergent. (**d**) Drug stimulated ATPase activities of wildtype EfrEF, the single mutants D41A^EfrE^ and D50A^EfrF^ and the corresponding double mutant (2xDtoA) reconstituted into proteoliposomes determined in the absence (basal activity) or in the presence of ethidium at the concentrations indicated. Data were normalized to the basal ATPase activity of the respective mutant. The error bars are standard deviations of technical triplicates. (**e**) Ethidium accumulation of *Lactococcus lactis* cells expressing wildtype EfrEF, the inactive Walker B mutant E515Q^EfrF^ or the extracellular gate mutants D41A^EfrE^ and D50A^EfrF^ or the corresponding double mutant (2xDtoA). (**f**) DEER analysis probing the extracellular and intracellular TMDs and the NBDs (same positions as in Fig. 2c). DEER traces were recorded in the presence of ATP-EDTA for the wildtype transporter and for TM287/288(2xDtoA). (**g**) Relative ATPase activities of the 2xDtoA mutant in the presence of increasing concentrations of Sb_TM#35, Nb_TM#1 and Nb_TM#2. A non-randomized sybody served as control. The data were fitted with a hyperbolic decay function to determine IC_50_ values as well as residual activities. The error bars are standard deviations of technical triplicates.

Using again ATP-EDTA to induce the IF to OF transition, DEER analyses revealed an equilibrium shift towards the OF state in the 2xDtoA mutant for all spin-labeled pairs (Fig. 4f and Fig. S9). The equilibrium shift was similar to that induced by Sb_TM#35. Hence, the aspartates at the extracellular gate constitute an energy barrier that needs to be overcome to switch to the OF state and influence the ATPase cycle in a long-ranged allosteric mechanism connecting the extracellular gate with the NBDs.

To probe the atomic details of the conformational dynamics underlying the IF-OF transition, we performed MD simulations of TM287/288 in a lipid bilayer starting from the IF crystal structure (PDB: 4Q4A), after docking a second ATP-Mg molecule into the consensus site^11^ and introducing the 2xDtoA mutations in the extracellular gate. As in our previous MD simulations of wildtype TM287/288 ^11^, we observed spontaneous large-scale conformational transitions from the IF conformation via an Occ state to an OF conformation; this complete transition was observed in 3 out of 20 independent 500 ns simulations (Fig. S10). Despite the limited statistics, the transition appears to be slightly more frequent than for the wildtype (6 out of 100 simulations^11^), in agreement with our experimental results and the notion that the polar contacts of the two aspartate residues increase the energy barrier of extracellular gate opening. Furthermore, 20 independent 400 ns simulations were carried out in which the OF structure reported in this work was used as starting structure, for both the wildtype and the 2xDtoA mutant. Although the sybody is not present in the simulations, the OF conformation with two ATP-Mg bound is very stable and merely fluctuates around the X-ray structure (Fig. S11).

Next, we introduced the 2xDtoA mutation into the heterodimeric ABC exporter EfrEF of *Enterococcus faecalis* (Fig. 4b)^15^. Ethidium-stimulated ATPase activity profiles of membrane reconstituted EfrEF were found to be strongly affected by the single DtoA mutations, and the ATPase activity of EfrEF containing the 2xDtoA could no longer be stimulated by the drug (Fig. 4d). Supporting this notion, the TM287/288(2xDtoA) mutant reconstituted in nanodiscs exhibited strongly diminished drug stimulation by Hoechst 33342 (Fig. S2a). Next, we expressed EfrEF wildtype and DtoA mutants in *Lactococcus lactis* and monitored ethidium uptake by fluorescence measurements (Fig. 4e). Wildtype EfrEF efficiently expels ethidium from the cell, resulting in a slow increase of ethidium accumulation that reaches a low steady-state level. EfrEF containing the EtoQ mutation in the NBDs served as negative control exhibiting high ethidium accumulation levels^16^. The single DtoA mutants D41A^EfrE^ and D50A^EfrF^ partially lost their capability of ethidium efflux. Interestingly, the accumulation curve of the 2xDtoA mutant does not reach a steady-state level within the time frame of the experiment. This observation suggests a transporter defect resulting in passive influx of ethidium into the cell mediated by EfrEF carrying the 2xDtoA mutation. In conclusion, the extracellular aspartates are important gate-keeper residues that are allosterically coupled to the NBDs and are required for substrate transport.

### Extracellular gate mutant and sybody are synergistic

Because both sybody binding and extracellular gate weakening shifted the conformational equilibrium towards the OF state, we reasoned that these effects are additive. Indeed, with an IC_50_ of 18.4 nM (Fig. 4g), inhibition of TM287/288 carrying the 2xDtoA mutation was found to be more pronounced than the inhibition of the wildtype transporter (IC_50_ = 66.1 nM) (Fig. 2b). In further agreement, the affinity of Sb_TM#35 towards the 2xDtoA mutant (K_D_ = 14 nM) was around eight times higher than towards the wildtype transporter (K_D_ = 110 nM) (Fig. S12, Table S2). Affinity was as well increased upon introduction of the EtoA or EtoQ mutation into the consensus site of the transporter (K_D_ = 66 nM) which as well exhibit a conformational equilibrium shift towards the OF state^9^ and was highest for the combined triple mutant (2xDtoA/EtoA) (K_D_ = 8 nM). An analogous pattern was observed for Nb_TM#1, which binds to the closed NBD dimer. The IC_50_ was substantially smaller when probing the 2xDtoA mutant (K_D_ = 16.8 nM) compared to wildtype TM287/288 (K_D_ = 460 nM) (Fig. 2b, Fig. 4g). This difference was again reflected by an affinity increase for the 2xDtoA mutant (37 nM) versus the wildtype transporter (K_D_ = 184 nM) and the highest affinity was observed for the triple mutant (K_D_ = 5 nM) (Fig. S12, Table S2). In aggregate, Sb_TM#35 and Nb_TM#1 bind to the opposite ends of the transporter but nevertheless exhibit a highly similar biophysical behavior and exclusively bind to and thereby trap the OF transporter.

### Probing the OF-IF conversion by state-specific binders

We finally tested whether the state-specific nanobodies could be used as probes in SPR to investigate the OF-IF transition of TM287/288. When TM287/288(EtoA) was charged with ATP-Mg and subsequently washed with buffer devoid of nucleotides, the maximal SPR binding signal for the OF state-specific binders Sb_TM#35 and Nb_TM#1 slowly decreased over a time window of several hours (Fig. 5a). Immobilized TM287/288 was stable within this time frame, because the maximal binding signal for the state-unspecific nanobody Nb_TM#2 only slightly decreased (Fig. 5a). Subsequently, we interrogated the OF-IF conversion using either ATP-Mg (hydrolyzing conditions) or ATP-EDTA (ATP binding without hydrolysis).

**Figure 5.**
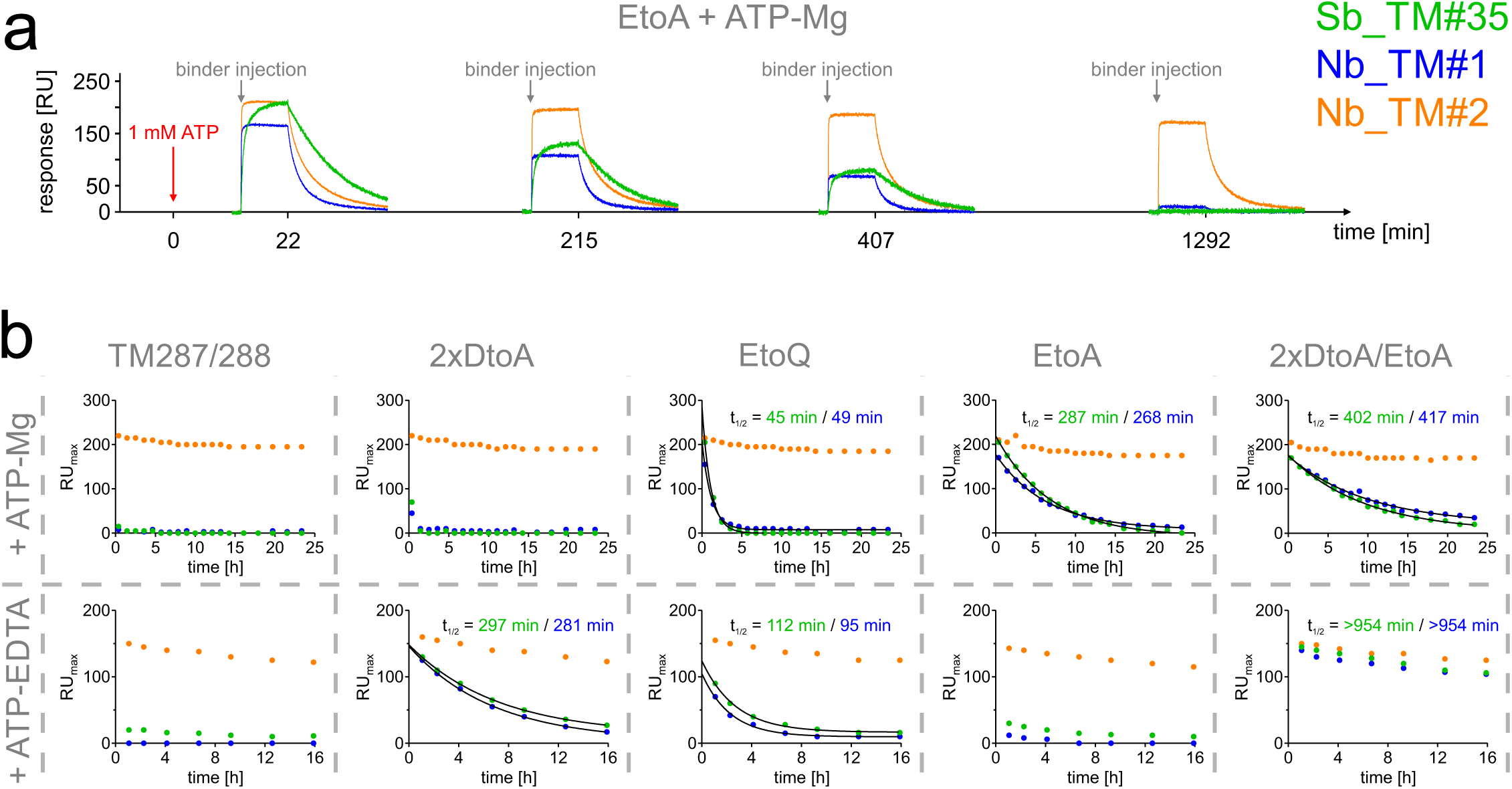
Probing the OF-IF transition using state-specific binders. (**a**) Exemplary raw data of SPR traces based on which conformational probing was detected. At time point zero, immobilized TM287/288(EtoA) was charged with 1 mM ATP (red arrow) and binders were injected (grey arrows) at saturating concentrations (Sb_TM#35, 1 µM; Nb_TM#1, 500 nM; Nb_TM#2, 100 nM) to obtain maximal response unit values (RU_max_) at the indicated time points. (**b**) RU_max_ values for wildtype and mutant TM287/288 were obtained as shown in (**a**) by charging the transporter with 1 mM ATP either in presence of Mg (upper panel) or EDTA (lower panel). For state-specific binders Sb_TM#35 and Nb_TM#1, data were fitted using a one phase decay function to determine half-life values (t_1/2_) of the OF state. State-unspecific Nb_TM#2 was used as a control.

In the case of wildtype TM287/288 loaded with ATP-Mg or ATP-EDTA, the state-specific binders were unable to recognize the transporter within the time resolution of the experiment, showing rapid conversion to the IF state. In contrast, the OF state was long-lived when ATP-Mg was occluded by the EtoQ or the EtoA mutant. The lifetime of the OF state as probed by the state-specific binders Sb_TM#35 or Nb_TM#1 was highly similar (t_1/2_ of 45 or 49 min for the EtoQ mutant and 287 or 268 min for the EtoA mutant, respectively). Strikingly, these values are in close agreement with the half-life of ATP hydrolyzed by these mutants, namely 32 min (EtoQ mutant) and 205 min (EtoA mutant) at 25°C. For the same mutants, the situation was inversed for ATP-EDTA. The EtoA mutant readily converted to the IF state akin to the wildtype transporter, whereas the OF state was very stable for the EtoQ mutant (t_1/2_ = 112 or 95 min, respectively). In this case, ATP cannot be hydrolyzed and we investigate NBD dissociation with bound ATP (but lacking the coordination by Mg^2+^). The glutamine in the Walker B position appears to stabilize ATP binding in the NBD sandwich dimer, whereas the canonical glutamate or an alanine at the same position promotes fast NBD dissociation. When the 2xDtoA mutation was introduced, the OF state was very long-lived in case of ATP-EDTA (t_1/2_ = 297 or 281 min, respectively). This suggests that weakening of the closed extracellular gate by the 2xDtoA mutation strongly impedes NBD dissociation, whereas NBD dissociation of the wildtype transporter is very fast under these experimental conditions. NBD opening is further slowed down if the 2xDtoA mutation is combined with the EtoA mutation for both ATP-Mg and ATP-EDTA.

From this dataset one can draw two major conclusions. First, ATP hydrolysis weakens the NBD dimer and is required to reset the transporter to the IF state under physiological conditions where ATP and Mg^2+^ are always present. And second, strong contacts at the extracellular gate are mandatory to exert a mechanical force onto the NBDs to facilitate fast NBD opening in order to reset the transport back to its IF state.

## DISCUSSION

In this work we unleashed the power of state-specific single domain antibodies obtained from alpacas and entirely *in vitro* from synthetic libraries to investigate a membrane transporter at the structural and functional level. The strategy of generating state-specific binders against type I ABC exporters has a long history going back to the 90’s of the last century, when the state-specific ABCB1 antibody UIC2 was identified^17^. A recent cryo-EM structure of UIC2 in complex with ABCB1 revealed that the antibody clamps the extracellular loops together, thereby preventing extracellular gate opening^18^. Molecular clamping of the closed extracellular gate was also achieved by a cyclic peptide raised against CmABCB1^19^. Further examples of binders that prevent the IF-OF conversion are nanobodies raised against ABCB1 and PglK, which both sterically clash with NBD closure^20, 21^. In contrast, our binders are specific for the OF state and consequently impede the OF-IF conversion.

Sybody binding to an extracellular wing of TM287/288 was essential to solve the OF structure. Both the degenerate and the consensus ATP binding site are fully closed and highly symmetric, but only the consensus site bears the catalytic dyad positioned to catalyze ATP hydrolysis^22^. This means that ATP hydrolysis of only one nucleotide is sufficient to initiate dissociation of the NBDs. In further support of this view, the closed NBD dimers features two possible P_i_ exit tunnels at the consensus site.

A comparison with other OF and Occ transporters revealed major conformational heterogeneity in the degree of extracellular gate opening, which has been discussed to play a potential role in squeezing out substrates or to prevent rebinding of substrates^3, 4, 10, 12, 23^. In this study, we uncover cross-talk between the extracellular gate and the ATPase cycle, a connection that has to the best of our knowledge not yet been investigated at the molecular level (Fig. 6). A sybody stabilizing the opened extracellular wing or mutations weakening the closed extracellular gate both shifted the conformational equilibrium towards the OF state. Previous experimental and computational studies have uncovered that NBD closure precedes extracellular gate opening during the IF-OF transition ^11, 24^, and DEER analyses have revealed that extracellular gate opening can be partial while NBD closure is complete ^9, 10, 25^. Hence, there seems to be an inbuilt mechanical principle that the extracellular gate is energetically costly to open. Conversely, using our state-specific nanobodies as conformational probes we were able to show that extracellular gate closure is coupled to the dissociation of the closed NBD dimer after substrate release and ATP hydrolysis. Our experiments on EfrEF demonstrated that a firmly sealed extracellular gate is in fact crucial for transporter function. Further, the 2xDtoA mutant had a strongly reduced ATPase activity and lost its capacity to be stimulated by drugs. This suggests that extracellular gate closure has become the rate-limiting step of the catalytic cycle of the 2xDtoA mutant. Hence, in this mutant the step of IF-OF conversion is no longer rate-limiting and consequently cannot be stimulated by drug binding to the inward-oriented high-affinity site (Fig. 6).

**Figure 6.**
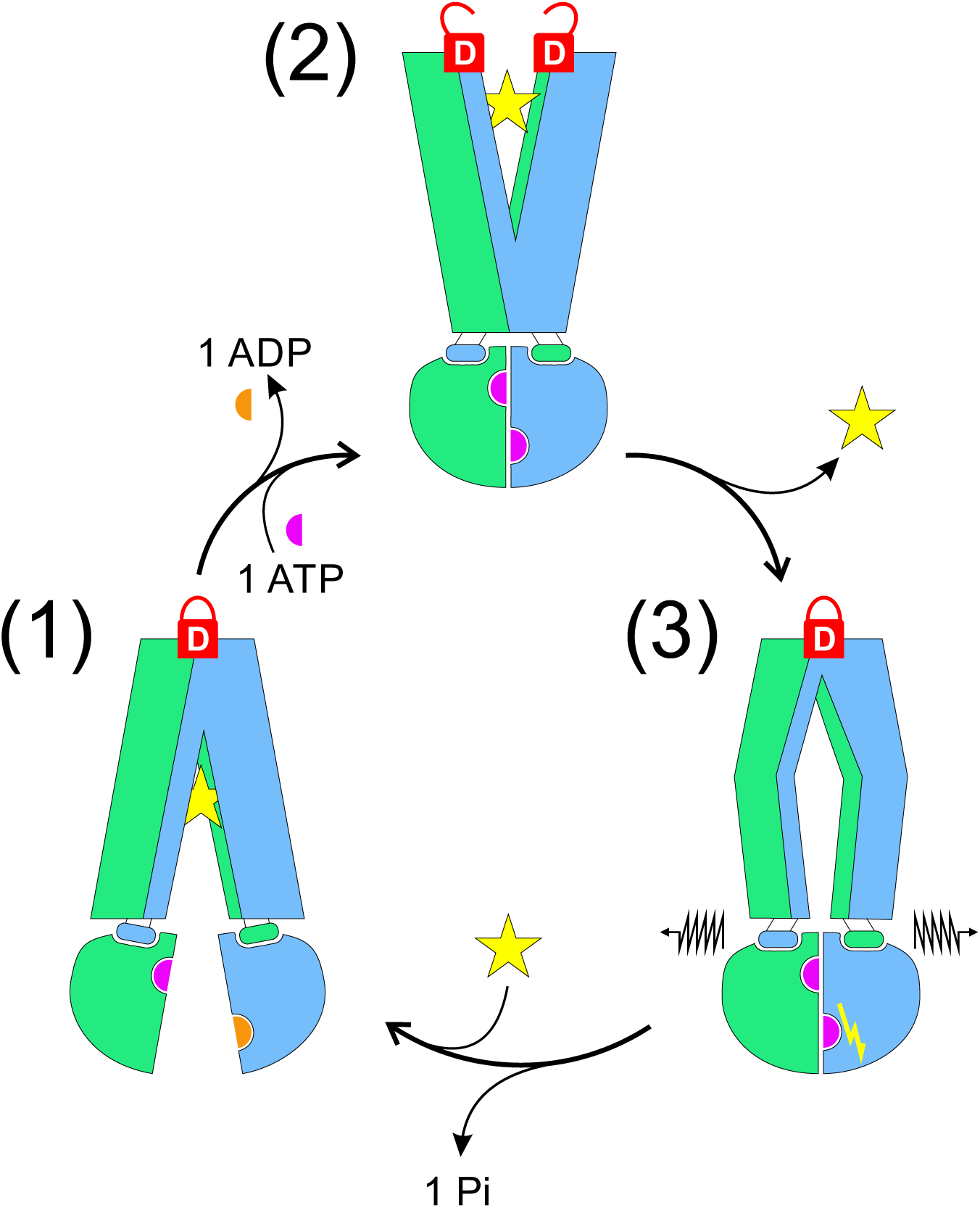
Role of the extracellular gate in the transport cycle of TM287/288. Substrate (yellow star) binds to the IF transporter (1) with high affinity, while the extracellular gate is sealed by two aspartates (closed D-lock). Binding and occlusion of two ATP-Mg molecules at the NBD interface drives the transition to the OF state (2). The extracellular gate opens (open D-lock) and substrate is released. The extracellular gate of the empty outward-oriented cavity closes (3) and thereby may trigger ATP hydrolysis at the consensus site. The mechanical force of the firmly sealed extracellular gate (closed D-lock) is required to dissociate the NBDs after ATP hydrolysis in order to reset the transporter to its IF state.

The ATP-bound OF state with completely closed NBDs has been referred to as the high-energy state of the transport cycle and in some instances it was proposed that ATP hydrolysis is needed to populate the OF state at all^10, 25^. In contrast, we and others have previously stipulated that ATP binding alone is sufficient for IF-OF conversion and substrate release^9, 11, 26^, in agreement with the ATP-switch model^27^. It should be noted that while the opened extracellular gate indeed represents a high-energy state, the opposite is in fact true for the closed NBD dimer. It adopts a low-energy state and a large energy input is required to dissociate the dimer^28^. This is certainly achieved in part by the hydrolysis of ATP^29^. Importantly, our results suggest that NBD dissociation also involves a mechanical component mediated by extracellular gate closure. A further possibility is the triggering of ATP hydrolysis as a result of extracellular gate closure. Although speculative and neither directly supported nor excluded by our data, such a mechanism would assure that the transporter only reverts to the IF state after substrate release.

Why and when ATP hydrolysis is required to achieve active transport is a recurrent debate in the ABC transporter field. Our data presented here and in previous studies^9, 11^ clearly suggest that ATP binding alone (in case of ATP-EDTA when ATP hydrolysis cannot occur) is sufficient for the IF-OF conversion and presumably the active transport of one substrate molecule. Directionality of transport is then achieved by an affinity switch of the substrate binding site, which inevitably undergoes drastic rearrangements as the TMDs switch from an IF to an OF conformation^12^. Nevertheless, it is unlikely that an ABC exporter can operate efficiently by binding and dissociation of ATP alone, because energy input is lacking and the molecular events would be driven by slow stochastic Brownian motions alone. To strongly populate the OF state under physiological conditions, ATP-Mg needs to be occluded at the consensus site of the closed NBD dimer^30^, a state that can be efficiently mimicked by the Walker B EtoQ or EtoA mutation^31^. As we show here with our binder probes, the resulting occluded ATP state traps the transporter in the OF state and prevents transporter cycling. Hence, ATP hydrolysis is needed to initiate dissociation of the closed NBD dimer. Provided the fact that ATPase activity returns to basal levels in the presence of high drug concentrations, it is reasonable to assume that the hydrolysis event happens after substrate release and is triggered by extracellular gate closure. Once ATP is hydrolyzed, the force exerted by the closed extracellular gate facilitates NBD dissociation. In summary, our results support the notion that ATP hydrolysis is required to drive the transport cycle at the resetting step from the OF to the IF state. We anticipate that our results provide a mechanistic framework to understand the functional role of the extracellular gate of type I ABC exporters and to explain the molecular underpinning of disease-causing mutations found in the extracellular region of medically important ABC exporters as recently investigated for MRP1 and CFTR^32, 33^.

## MATERIALS AND METHODS

### Expression and purification

TM287/288 was expressed in *E. coli* MC1061 and purified for crystallization and functional assays as previously described^7^. For some crystallization experiments, β-DDM was exchanged by 0.3% (w/v) n-decyl-β-D-maltoside (β-DM, Glycon) during the affinity chromatography step on a Ni-NTA Superflow (Qiagen) gravity flow column. Biotinylated TM287/288 was prepared as described^14^. Purified TM287/288 variants were flash frozen in liquid nitrogen and stored at -80°C ready to use. For crystallization, TM287/288 was always prepared freshly. EfrEF was expressed in *L. lactis* NZ9000*δlmrAδlmrCD*, purified and reconstituted in proteoliposomes as described previously^15^. Nanobodies/sybodies were either expressed from pSb_init (addgene: #110100) for biochemical experiments or sub-cloned into pBXNPMH3 (addgene: #110099) by FX cloning for the production of tag-free nanobodies/sybodies for crystallization and DEER experiments^14, 34^. Nanobodies/sybodies were expressed and purified as described previously^14^ and stored at -80°C.

### Mutagenesis

To weaken the extracellular gate, two conserved aspartates were replaced by alanines in various TM287/288 variants resulting in TM287/288(2xDtoA). D41A^TM287^ was introduced using the primers TM287_D41A_FW (GGC ACG TAT TGT CGC CGA AGG AAT CG C) and TM287_D41A_RV (GCG ATT CCT TCG GCG ACA ATA CGT GCC). D65A^TM288^ was introduced using the primers TM288_D65A_FW (CAT AGG AAA AAC GAT CGC TGT TGT CTT CG) and TM288_D65A_RV (CGA AGA CAA CAG CGA TCG TTT TTC CTA TG). In order to render TM287/288 catalytically inactive, E517A^TM288^ was introduced in wildtype TM287/288 and TM287/288(2xDtoA) using the primers TM288_E517A_FW (CCT GAT ACT GGA CGC AGC CAC CAG CAA C) and TM288_E517A_RV (GTT GCT GGT GGC TGC GTC CAG TAT CAG G). Mutation D41A^EfrE^ was introduced using the primers EfrE_D41A_FW (CAA GTT GAT TGC TGT GGG CAT CG) and EfrE_D41A_RV (CGA TGC CCA CAG CAA TCA ACT TG). D50A^EfrF^ was generated using the primers EfrF_D50A_FW (CAA TCG GGA TTG CTA ACC TCT TAG AAG C) and EfrF_D50A_RV (GCT TCT AAG AGG TTA GCA ATC CCG ATT G). In order to cross-link the transporter at the tetrahelix bundle in the OF state, L200C^TM287^ was introduced in cys-less TM287/288 (described in ^7^) using the primers TM287_L200C_FW (GAG AAA ATC TCT GCG GTG TCA GGG TAG TGA G) and TM287_L200C_RV (CTC ACT ACC CTG ACA CCG CAG AGA TTT TCT C) and combined with the S224C^TM288^ mutation using the primers TM288_S224C_FW (CAT AGA AGA AGA CAT CTG CGG CCT CAC TGT G) and TM288_S224C_RV (CAC AGT GAG GCC GCA GAT GTC TTC TTC TAT G).

For spin-labeling of the sybody, S71C was introduced in Sb_TM#35 using the primers Sb_TM#35_S71C_FW (CAC GGT GTG CCT GGA CAA CG) and Sb_TM#35_S71C_RV (CGT TGT CCA GGC ACA CCG TG).

For spin-labeling of TM287/288, three new cysteines were introduced in cys-less TM287/288 (called wildtype TM287/288 for simplicity). S271C^TM287^ was generated using the primers TM287_S271C_FW (CAG ATG GAG ATA GGA TGC ATC ATG GCA TAC) and TM287_S271C_RV (GTA TGC CAT GAT GCA TCC TAT CTC CAT CTG) as a single mutant or in combination with K54C^TM287^ which was generated using the primers TM287_K54C_FW (CTT TTC TCT GGT TTT GTG TAC AGG GAT CCT CAT G) and TM287_K54C_RV (CAT GAG GAT CCC TGT ACA CAA AAC CAG AGA AAA G). K54C^TM287^ was also prepared as a single mutant or in combination with I290C^TM288^ introduced with the primers TM288_I290C_FW (CGC CTT GAA AGA CTG TAT CAC GGT GGG) and TM288_I290C_RV (CCC ACC GTG ATA CAG TCT TTC AAG GCG).

### Crystallization

For crystallization of TM287/288(EtoA) in complex with Sb_TM#35, freshly purified transporter in 0.3% (w/v) β-DM was concentrated to about 12 mg/ml using an Amicon Ultra-4 concentrator unit (50 kDa MWCO) and purified Sb_TM#35 (stored at -80°C) was added in a 1.1-fold molar excess (final complex concentration of 10 mg/ml). Complexes were pre-incubated with 2.5 mM adenosine 5′-(3-thiotriphosphate) (ATPγS, Sigma, A1388), 3mM MgCl_2_ for 5-6 days at 20°C (without this incubation step, the crystals did not diffract to high resolution), before crystals were grown by the vapour diffusion method in sitting drops (1:1 protein to reservoir ratio) at 20°C in 0.1 M Na-acetate pH 4.6, 0.035 M NaCl and 11.5% (w/v) PEG6000. Crystals appeared within 1-3 days and were fished immediately. Crystals were cryo-protected in 0.1 M Na-acetate pH 4.6, 0.04 M NaCl and 15% (w/v) PEG6000 additionally containing 20 mM Tris-HCl pH 7.5, 150 mM NaCl, 0.3% (w/v) β-DM, 3 mM MgCl_2_, 1.25 mM ATPγS and 25% (v/v) ethylenglycol and flash-frozen in liquid nitrogen.

For crystallization of TM287/288(2xDtoA/EtoA) in complex with Nb_TM#1, purified Nb_TM#1 (stored at -80°C) was added to the transporter purified in 0.3% (w/v) β-DM in a 1.2-fold molar excess prior to size-exclusion chromatography. After short incubation on ice, the complex was separated from excess nanobodies by size-exclusion chromatography using a Superdex 200 Increase 10/300 GL (GE Healthcare) column equilibrated in 20 mM Tris-HCl pH 7.5, 150 mM NaCl and 0.3% (w/v) β-DM. The transporter/nanobody complexes were concentrated to 10 mg/ml using an Amicon Ultra-4 concentrator unit (50 kDa MWCO) and incubated with 5 mM ATPγS and 3 mM MgCl_2_ for 15 min on ice. Crystals were grown by the vapour diffusion method in sitting drops (1:1 protein to reservoir ratio) at 20°C in 0.05 M Glycine pH 9.5, 0.225 M NaCl and 21% (v/v) PEG550MME. Crystals appeared within 2-3 days and were grown for another 3 weeks. Crystals were cryo-protected in reservoir solution containing 10% (v/v) PEG400 and flash-frozen in liquid nitrogen.

For crystallization of TM287/288(2xDtoA/EtoA) in complex with Nb_TM#2, purified Nb_TM#2 (stored at -80°C) was added to the cleaved transporter purified in 0.03% (w/v) β-DDM in a 1.2-fold molar excess prior to size-exclusion chromatography. After short incubation on ice, the complex was separated from excess nanobodies by size-exclusion chromatography using a Superdex 200 Increase 10/300 GL (GE Healthcare) column equilibrated in 20 mM Tris-HCl pH 7.5, 150 mM NaCl and 0.03% (w/v) β-DDM. The transporter/nanobody complexes were concentrated to 10 mg/ml using an Amicon Ultra-4 concentrator unit (50 kDa MWCO) and incubated with 2.5 mM ATPγS and 3 mM MgCl_2_ for 15 min on ice. Crystals were grown by the vapour diffusion method in sitting drops (1:1 protein to reservoir ratio) at 20°C in 0.1 M Tris-HCl pH 8.5, 0.1 M NaCl and 30% (v/v) PEG400. Crystals appeared within 2-3 days and were grown for another 2-3 weeks. Crystals were flash-frozen in liquid nitrogen without further cryo-protection.

### Data collection and structure determination

Diffraction data were collected with a wavelength of 1.0 Å at 100 K at the beamlines X06DA and X06SA at the Swiss Light Source (SLS, Villigen, Switzerland). Diffraction data were processed with the program XDS^35^ and truncated using the Diffraction Anisotropy Server with default settings^36^ due to strong or severe anisotropy, what lead to improved electron density maps (Table S1, Fig. S13).

The TM287/288(EtoA) – Sb_TM#35 – ATPγS-Mg complex structure was solved by molecular replacement in Phaser^37^ using a modified homology model based on Sav1866 (PDB: 2HYD). The crystals belong to the space group P2_1_ containing two TM287/288 heterodimers and two sybodies per asymmetric unit. After a few cycles of model building in Coot^38^ and refinement in Buster (www.globalphasing.com), a poly-alanine model of a nanobody (PDB: 1ZVH) was manually placed into additional electron density. Multiple iterations of model building in Coot and TLS refinement in Buster resulted in a final model with good geometry (Ramachandran favored/outliers: 96.57% / 0.16%) (Table S1). Chains A (TM287), B (TM288) and E (Sb_TM#35) were used for structural analysis and figures.

The TM287/288(2xDtoA/EtoA) – Nb_TM#1 – ATPγS-Mg complex structure was solved by molecular replacement in Phaser using the TM287/288(EtoA) – Sb_TM#35 structure without the sybody. The crystals belong to the space group P2_1_ containing two TM287/288 heterodimers and two nanobodies per asymmetric unit. After some cycles of model building in Coot and refinement in Buster, Phaser was used to place the missing nanobody using a homology model based on PDB entry 5OCL. Multiple iterations of model building in Coot and TLS refinement in Buster resulted in a final model with good geometry (Ramachandran favored/outliers: 94.84% / 0.44%) (Table S1). Chain A (TM287), B (TM288) and E (Nb_TM#1) were used for figures.

The TM287/288(2xDtoA/EtoA) – Nb_TM#2 – ATPγS-Mg complex structure was solved by molecular replacement in Phaser using the TM287/288(EtoA) – Sb_TM#35 structure without the sybody. The crystals belong to the space group P1 containing two TM287/288 heterodimers and two nanobodies per asymmetric unit. After several cycles of model building in Coot and refinement in Buster, Phaser was used to find the missing nanobody using a poly-alanine homology model based on PDB entry 4LAJ. Multiple iterations of model building in Coot and TLS refinement in Buster resulted in a final model at 4.2 Å resolution (Ramachandran favored/outliers: 94.09% / 0.83%) (Table S1). Chain A (TM287), B (TM288) and E (Nb_TM#2) were used for figures. Molecular graphics and analyses were performed in Pymol or with UCSF Chimera^39^.

### ATPase assays

ATPase activities were measured by detecting liberated phosphate using molybdate/malachite green detection as described previously^8^. ATPase activity measurements with detergent-purified protein were carried out in ATPase buffer consisting of 20 mM Tris-HCl pH 7.5, 150 mM NaCl, 10 mM MgSO_4_ containing 0.03% (w/v) β-DDM. Activity measurements of TM287/288 reconstituted in nanodiscs were performed in the same buffer lacking detergent. Measurements of EfrEF reconstituted in proteoliposomes were carried out in 50 mM HEPES pH 7.0 and 10 mM MgSO_4_.

Relative ATPase activities of TM287/288(D41A^TM287^), TM287/288(D65A^TM288^) and TM287/288(2xDtoA) compared to wildtype TM287/288 were measured at 25°C for 15 min in the presence of 500 µM ATP. 40 nM wildtype TM287/288, 80 nM single DtoA TM287/288 variants and 160 nM TM287/288(2xDtoA) were used and the respective concentration of TM287/288(EtoQ) for background subtraction.

The relative ATPase activity stimulations of EfrEF variants in proteoliposomes by ethidium were determined at 30°C for 15 min in the presence of 1 mM ATP. The amount of reconstituted EfrEF variants was determined by quantitative SDS-PAGE. 4 nM wildtype EfrEF, 23 nM single DtoA EfrEF variants and 15 nM EfrEF(2xDtoA) were used and buffer controls were taken for background subtraction.

Inhibition of ATPase activities of wildtype TM287/288 and TM287/288(2xDtoA) in detergent by binders was determined in the presence of 500 µM ATP at 25°C for 30 min or 60 min, respectively. 8 nM wildtype TM287/288 as well as TM287/288(2xDtoA) were used, which is more than 2-fold less than the lowest binder concentration (20 nM). For background subtraction equal amounts of TM287/288(EtoQ) were used. To obtain IC_50_ values, the inhibition data were fitted with a hyperbolic decay curve with the following function (SigmaPlot):

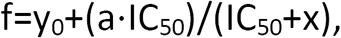

in which f corresponds to the ATPase activity at the respective binder concentration concentration divided by the ATPase activity in the absence of inhibitor normalized to 100%, y_0_ corresponds to the residual activity at infinite binder concentration, a corresponds to the maximal degree of inhibition (a+y_0_=100%) and x corresponds to the binder concentration.

Stimulated ATPase activities of wildtype TM287/288 and TM287/288(2xDtoA) in nanodiscs were determined at 37°C for 30 min in the presence of 500 µM ATP and varying Hoechst 33342 concentrations. 50 nM wildtype TM287/288 as well as TM287/288(2xDtoA) were used to determine relative stimulations compared to basal ATPase activities using buffer for background subtraction.

Hoechst 33342 stimulated ATPase activity inhibition in nanodiscs was determined at 37°C for 30 min in the presence of 500 µM ATP and 50 µM Hoechst 33342. 10 nM wildtype TM287/288 and 50 nM TM287/288(2xDtoA) in nanodiscs were used to determine ATPase activities in the presence or absence of 10 µM binders using empty nanodiscs for background subtraction.

### BMOE cross-linking

To raise alpaca nanobodies specifically recognizing the OF state of TM287/288, the transporter was cross-linked at the tetra-helix bundle, which forms when the transporter adopts the OF state^13^. Two cysteines were introduced in the cys-less TM287/288 variant at positions L200C^TM287^ and S224C^TM288^ by site-directed mutagenesis. On top, the 2xDtoA mutations were introduced in TM287/288_cl_L200C^TM287^/S224C^TM288^. Since the two cysteines are too far apart to form a disulfide bond, the maleimide cross-linker BMOE (bismaleimidoethane, Thermo Scientific™, #22323) with a length of 8 Å was used. TM287/288_cl_L200C^TM287^/S224C^TM288^ with or without the 2xDtoA mutations was expressed as described above. Membranes were solubilized and purified by Ni-NTA affinity chromatography in presence of 1 mM dithiothreitol (DTT) in 0.03% (w/v) β-DDM. Buffer was exchanged and DTT removed by size-exclusion chromatography using a Superdex 200 Increase 10/300 GL (GE Healthcare) column equilibrated in PBS pH 7.4 and 0.03% (w/v) β-DDM at 4°C and concentrated to 50 µM with an Amicon Ultra-4 concentrator unit with a MWCO of 50 kDa. 10 mM ATP, 3 mM MgCl_2_ and a 5-fold molar excess of BMOE over transporter were added and the cross-linking mixture incubated for 3 h at 30°C. The mixture was diluted 5-fold and incubated with 3C protease (1:10 w/w) overnight at 4°C. The cross-linked and Tag-free transporter was reloaded on a Ni-NTA gravity flow column equilibrated with 20 mM Tris-HCl pH 7.5, 150 mM NaCl, 40 mM imidazole and 0.03% (w/v) β-DDM. The sample was concentrated with Amicon Ultra-4 concentrator units with a MWCO of 50 kDa, 10% (v/v) glycerol was added and aliquots were flash frozen in liquid nitrogen and stored at -80°C ready for size-exclusion chromatography. Cross-linked TM287/288_cl_L200C^TM287^/S224C^TM288^ with or without the 2xDtoA mutations was polished by size-exclusion chromatography using a Superdex 200 Increase 10/300 GL (GE Healthcare) column equilibrated in 20 mM Tris-HCl pH 7.5, 150 mM NaCl and 0.03% (w/v) β-DDM at 4°C and concentrated to 1 mg/ml with an Amicon Ultra-4 concentrator unit with a MWCO of 50 kDa and immediately used for alpaca immunizations.

### Nanobody and sybody selections

For the selection of OF state-specific nanobodies, an alpaca was immunized with subcutaneous injections four times in two week intervals, each time with 200 µg purified cross-linked TM287/288(2xDtoA)_cl_L200C^TM287^/S224C^TM288^ in 20 mM Tris-HCl pH 7.5, 150 mM NaCl and 0.03% (w/v) β-DDM. Blood was collected two weeks after the last injection for the preparation of the lymphocyte RNA, which was then used to generate cDNA by RT-PCR to amplify the VHH/nanobody repertoire. Phage libraries were generated as described^40^ and two rounds of phage display were performed against TM287/288(2xDtoA/EtoA) solubilized in β-DDM in the presence of 2 mM ATP as described^14^. After the final phage display selection round, 91.9-fold enrichment was determined by qPCR using AcrB as background. The enriched nanobody library was sub-cloned into pSb_init by FX-cloning and 94 single clones were analyzed by ELISA in the presence of 1 mM ATP as described^14^. The 27 positive ELISA hits were Sanger sequenced and grouped in three families according to their CDR3 length and sequence (among them was Nb_TM#1). In another selection cross-linked TM287/288_cl_L200C^TM287^/S224C^TM288^ was used for alpaca immunizations resulting in 51 positive ELISA hits, of which 24 were Sanger sequenced and grouped into four binder families (among them was Nb_TM#2). The selection of sybodies against the OF state of TM287/288 was described in detail in a previous study^14^.

### Spin labeling for DEER

Spin-labeled TM287/288 variants were expressed, purified and spin labeled with MTSL [(1-oxyl-2,2,5,5-tetramethyl-δ3-pyrroline-3-methyl)methanethiosulfonate, Toronto Research] as described elsewhere^8^. The pairs 131^TM288^/248^TM288^ and 460^TM287^/363^TM288^ were already used in previous studies^8, 9^, whereas the extracellular spin-label pairs 54^TM287^/271^TM287^ and 54^TM287^/290^TM288^ were constructed as part of this study. The samples were concentrated to 30-50 µM with Amicon Ultra-4 concentrator units with a MWCO of 50 kDa, flash-frozen in liquid nitrogen and stored at -80°C.

For site specific spin-labeling of Sb_TM#35, a single cysteine was introduced in the framework of the sybody at position 71 by site-directed mutagenesis. Sb_TM#35_S71C was expressed from pBXPHM3 as described above. Cells were harvested and resuspended in PBS pH 7.4 and 2mM DTT supplemented with DNase (Sigma) and disrupted with an M-110P Microfluidizer^®^ (Microfluidics™). The supernatant, supplemented with 20 mM imidazole, was loaded on a Ni-NTA gravity flow column, washed with 20 column volumes PBS pH 7.4, 50 mM imidazole and 2 mM DTT and eluted with 4 column volumes PBS pH 7.4, 300 mM imidazole and 2 mM DTT.

In a next step Sb_TM#35_S71C fused to His-tagged MBP was incubated with 3C protease (1:10 w/w), while dialyzing against PBS pH 7.4 and 2 mM DTT overnight at room temperature. Cleaved sybody was reloaded on a Ni-NTA gravity flow column and eluted with 3 column volumes PBS pH 7.4, 40 mM imidazole and 2 mM DTT, followed by size exclusion chromatography using a Sepax-SRT10C SEC-300 (Sepax Technologies) column equilibrated with PBS pH 7.4 and 2 mM DTT. Peak fractions were collected and DTT removed using a PD-10 (GE Healthcare, 17-0851-1) column equilibrated with degassed PBS pH 7.0 at 4°C. To avoid DTT take-over, the sybodies were eluted with 3.2 ml degassed PBS pH 7.0, instead of the 3.5 ml suggested by the manufacturer. The elution was concentrated to 2.5 ml using an Amicon concentrator unit with a 3 kDa MWCO. 5-fold molar excess MTSL was added to the sample and incubated for 1 h on ice, a condition which was previously reported to prevent labeling of the buried cysteines that form the universally conserved disulfide bond of nanobodies^41^. Free label was removed and buffer exchanged using a PD-10 column (GE Healthcare, 17-0851-1) equilibrated with 20 mM Tris-HCl pH 7.5 and 150 mM NaCl. In order to avoid MTSL take-over, the sybodies were eluted with 3.2 ml 20 mM Tris-HCl pH 7.5 and 150 mM NaCl. Spin-labeled nanobodies were concentrated to the desired concentration with an Amicon Ultra-4 concentrator unit with a 3 kDa MWCO and flash-frozen in liquid nitrogen and stored at -80°C. Site-specific labeling was confirmed and quantified by mass spectrometry.

### DEER measurements

The labeling efficiency of the double cysteine mutants of the transporters solubilized in detergent was measured by comparing the second integral of the spectra detected at 25°C using an X-band Miniscope 400 EPR spectrometer (Magnettech by Freiberg Instruments) with that of a standard TEMPOL solution in water. The calculated spin labeling efficiencies of the twelve mutants ranged between 80 and 90%. For DEER measurements, 10% (v/v) D_8_-glycerol was added as cryoprotectant. The range of final transporter concentrations was 15 to 25 µM. 40 µL of sample was loaded in quartz tubes with 3 mm outer diameter. The ATP-EDTA sample contained 2.5 mM ATP and 2.5 mM ethylenediaminetetraacetate (EDTA) to completely inhibit ATP hydrolysis; samples were incubated at 25°C for 10 minutes and snap-frozen in liquid nitrogen. For vanadate trapping, samples were incubated with 5 mM sodium orthovanadate, 2.5 mM ATP and 2.5 mM MgCl_2_ for 3 min at 50°C and snap-frozen in liquid nitrogen. The unlabeled Sb_TM#35 was added to the TM287/288 in ∼1.3:1 stoichiometric ratio. To measure sybody-transporter distances, Sb_TM#35 spin-labeled at position 71 was added in a 0.5:1 stoichiometric ratio to the singly-labeled TM287/288 mutants (54^TM287^ and 271^TM287^) in the presence of ATP-EDTA.

Double electron-electron resonance (DEER) measurements were performed at 50 K on a Bruker ELEXSYS E580Q-AWG (arbitrary waveform generator) dedicated pulse Q-band spectrometer equipped with a 150 W TWT amplifier. A 4-pulse DEER sequence with Gaussian, non-selective observer and pump pulses of 32 or 34 ns length (corresponding to 14 or 16 ns FWHM) with 100 MHz frequency separation was used. Due to the coherent nature of the AWG generated pulses, a four-step phase cycling (0 - π/2 - π - 3/2π) of the pump pulse was performed together with 0 - π phase cycling of the observer pulses to remove unwanted effects of running echoes from the DEER trace. The evaluation of the DEER data was performed using DeerAnalysis2015 ^42^. The background of the primary DEER traces was corrected using stretched exponential functions with homogeneous dimensions of 1.8 to 3 for different samples. A model-free Tikhonov regularization was used to extract distance distributions from the background corrected form factors. The data of the apo and ATP-EDTA states shown for the pairs 131^TM288^/248^TM288^ and 460^TM287^/363^TM288^ in the absence of sybody are reproducible with respect to those previously published^9^. Interspin distance simulations were performed with the software MMM2015 using the MTSL ambient temperature library^43, 44^.

### Transport assay and reconstitution in proteoliposomes

Fluorescent dye transport with EfrEF in *L. lactis* was performed exactly as described previously^15, 16^. EfrEF purification and reconstitution into proteoliposomes was done exactly as described before^15^.

### Surface plasmon resonance

Binding affinities were determined by surface plasmon resonance (SPR) at 25°C using a ProteOn™ XPR36 Protein Interaction Array System (Biorad). Biotinylated TM287/288 variants were immobilized on ProteOn™ NLC Sensor Chips at a density of 2000 RU. Nanobodies and sybodies expressed in pSb_init were gel-filtrated in 20 mM Tris-HCl pH 7.5 and 150 mM NaCl, and the SPR measurements were carried out in the same buffer containing 0.015% (w/v) β-DDM and either 1 mM MgCl_2_ or 1 mM MgCl_2_ and 0.5 mM ATP to measure binding affinities in the absence or presence of ATP, respectively. Every measurement was done once and the data fitted with a 1:1 interaction model using the BioRad Proteon Analysis Software. In order to determine the half-lives of the outward-facing, ATP-bound state of the different TM287/288 variants, all five biotinylated variants were immobilized on a ProteOn™ NLC Sensor Chips at a density of 3000 RU. The experiment was conducted in 20 mM Tris-HCl pH 7.5, 150 mM NaCl, 0.015% (w/v) β-DDM and 1 mM MgCl_2_ or 2.5 mM EDTA at 25°C at a flow-rate of 30 µl/min. In order to charge the transporter variants with ATP-Mg or ATP-EDTA, buffer containing 1 mM ATP together with either 1 mM MgCl_2_ or 2.5 mM EDTA was injected at the beginning of the experiment.

### Nanodisc preparation

Membrane scaffold protein MSP1D1E3 was sub-cloned from pINITIAL (provided by Prof. Raimund Dutzler) into pBXNH3 (addgene: Plasmid #47067) by FX cloning^34^. Freshly transformed MC1061 *E. coli* cells were grown in Terrific Broth (TB) medium supplemented 100 µg/ml ampicillin to an OD_600_ of 1.0 – 1.5 at 37°C and expression was induced by the addition of 0.0017% (w/v) L-arabinose overnight at 22°C. Cells were harvested, resuspended in lysis buffer (20 mM Na-phosphate pH 7.4, 1% (v/v) Triton X-100 and 1 mM PMSF) and disrupted with a M-110P Microfluidizer^®^ (Microfluidics™). Cell debris were pelleted at 8000 g for 30 min at 4°C and the supernatant was loaded on a Ni-NTA gravity flow column equilibrated with lysis buffer, washed with 10 column volumes buffer 1 (40 mM Tris-HCl pH 8.0, 0.3 M NaCl and 1% (v/v) Triton X-100), 10 column volumes buffer 2 (buffer 1 + 50 mM sodium cholate), 10 column volumes buffer A (40 mM Tris-HCl pH 8.0 and 0.3 M NaCl), 10 column volumes buffer A containing 20 mM imidazole and finally eluted with 4 column volumes buffer A containing 300 mM imidazole. 3C protease was added (1:10 w/w) and the sample dialyzed overnight at 4°C against 20 mM Tris-HCl pH 7.5, 150 mM NaCl and 0.5 mM K-EDTA. The next day, 2 mM MgCl_2_ were added to the sample, which was then reloaded on a Ni-NTA gravity flow column equilibrated with 20 mM Tris-HCl pH 7.5, 150 mM NaCl and 20 mM imidazole and eluted with 2 column volumes using the same buffer. Buffer was exchanged by dialyzing three times against Tris-HCl pH 7.5 and 0.5 mM K-EDTA for 1.5 h at room temperature. The purified membrane scaffold protein was concentrated to 60 mg/ml using an Amicon Ultra-4 concentrator unit with a 10 kDa MWCO, flash-frozen in liquid nitrogen and stored at -80°C.

*E. coli* polar lipids (*E. coli* polar lipid extract, Avanti, 100600C) were mixed 3:1 (w/w) with L-α-phosphatidylcholine from egg yolk (Sigma, P3556) and chloroform was evaporated. Dried lipids were dissolved in 20 mM HEPES pH 8.0, 0.5 mM K-EDTA, 100 mM NaCl and 100 mM cholate to a final concentration of 38 mg/ml (50 mM), and filtered using a 0.22 µM filter. Lipids ready for nanodisc reconstitutions were stored at -80°C.

Purified, biotinylated TM287/288 variants solubilized in β-DDM were reconstituted into nanodiscs using a 240:8:1 molar ratio of lipids:MSP1D1E3:TM287/288. In order to spare membrane protein, the ideal lipid:MSP1D1E3 ratio was determined beforehand by reconstituting empty nanodiscs. Different ratios were tested, ranging from 25:1 to 40:1 (lipid:MSP1D1E3). Empty nanodiscs were loaded on a Superdex 200 Increase 10/300 GL (GE Healthcare) to separate empty nanodiscs from monomeric MSP1D1E3 and aggregates. From the elution profile the optimal lipid:MSP1D1E3 ratio of 30:1 was determined. The final cholate concentration in the reconstitution mixture was adjusted to 30 mM. The mixture was incubated at 25°C for 20 min while rocking at 650 rpm. Then, 200 mg bio-beads were added to 200 µl reconstitution mixture, which was incubated overnight at 4°C while rocking at 1000 rpm. Bio-beads were removed using 0.1 µm PVDF spin-filters. Full nanodiscs were separated from empty nanodiscs by gel filtration using a Superdex 200 Increase 10/300 GL (GE Healthcare) equilibrated with 20 mM Tris-HCl pH 7.5 and 150 mM NaCl. TM287/288 variants reconstituted in nanodiscs were either immediately used or flash-frozen in liquid nitrogen and stored at -80°C.

### MD simulations

The simulation setup and parameters are equivalent to our previous work^11^. In brief, all-atom MD simulations of TM287/288 embedded in an explicitly solvated POPC bilayer were carried out with GROMACS^45^. For the simulations initiated from the IF structure (PDB: 4Q4A), ATP-Mg was docked into the consensus site and D41^TM287^ and D65^TM288^ were replaced by alanines. We conducted 20 individual simulations of length 500 ns each (i.e., 10 microseconds in total) at 375 K. In addition, the OF state was simulated starting from the crystal structure shown in Fig. 1a after removing the sybody and replacing ATP-g-S by ATP. Furthermore, the same set of simulations was carried out for the ATP-bound 2xDtoA OF structure (another 8 microseconds in total).

## ACKNOWLEDGEMENTS

We thank all members of the Seeger lab for stimulating discussions. We acknowledge Beat Blattmann and Céline Stutz-Ducommun of the Protein Crystallization Center UZH for performing the crystallization screening, and the staff of the SLS beamlines X06SA and X06DA for their support during X-ray data collection. EB would like to thank G. Jeschke (ETH Zurich) for providing the Q-band resonator. The Steinbuch Centre for Computing (SCC) in Karlsruhe/Germany provided computational resources. The Institute of Medical Microbiology and the University of Zurich are acknowledged for financial support. MK thanks Natural Sciences and Engineering Research Council of Canada (NSERC) and the Canada Research Chairs Program for financial support. This work was funded by a SNF Professorship of the Swiss National Science Foundation (PP00P3_144823, to MAS) and by the Deutsche Forschungsgemeinschaft (DFG) through an Emmy Noether grant to LVS (SCHA 1574/3-1), Cluster of Excellence RESOLV (EXC 1069), and research grant to EB (BO 3000/ 1-2 and INST 130/972-1 FUGG).

## AUTHOR CONTRIBUTIONS

CAJH, EB and MAS conceived the study. CAJH, IZ, PE and SS selected nanobodies and sybodies against TM287/288. CAJH purified and crystallized the protein complexes and solved their structure. LMH conducted the functional and biochemical experiments with EfrEF. CAJH and LMH conducted all functional experiments with TM287/288. CAJH cloned, purified and labeled all samples for DEER analyzes. MHT conducted most of the DEER analyzes and simulations and discussed the results with EB. SK conducted the sybody-transporter DEER experiments and discussed them with EB. HG carried out MD simulations and analyzed and interpreted them together with MK and LVS. CAJH, MHT, LMH, HG and MAS created figures. CAJH, LVS, EB and MAS wrote the manuscript and all authors edited the manuscript.

## FIGURE LEGENDS

**Figure S1.**
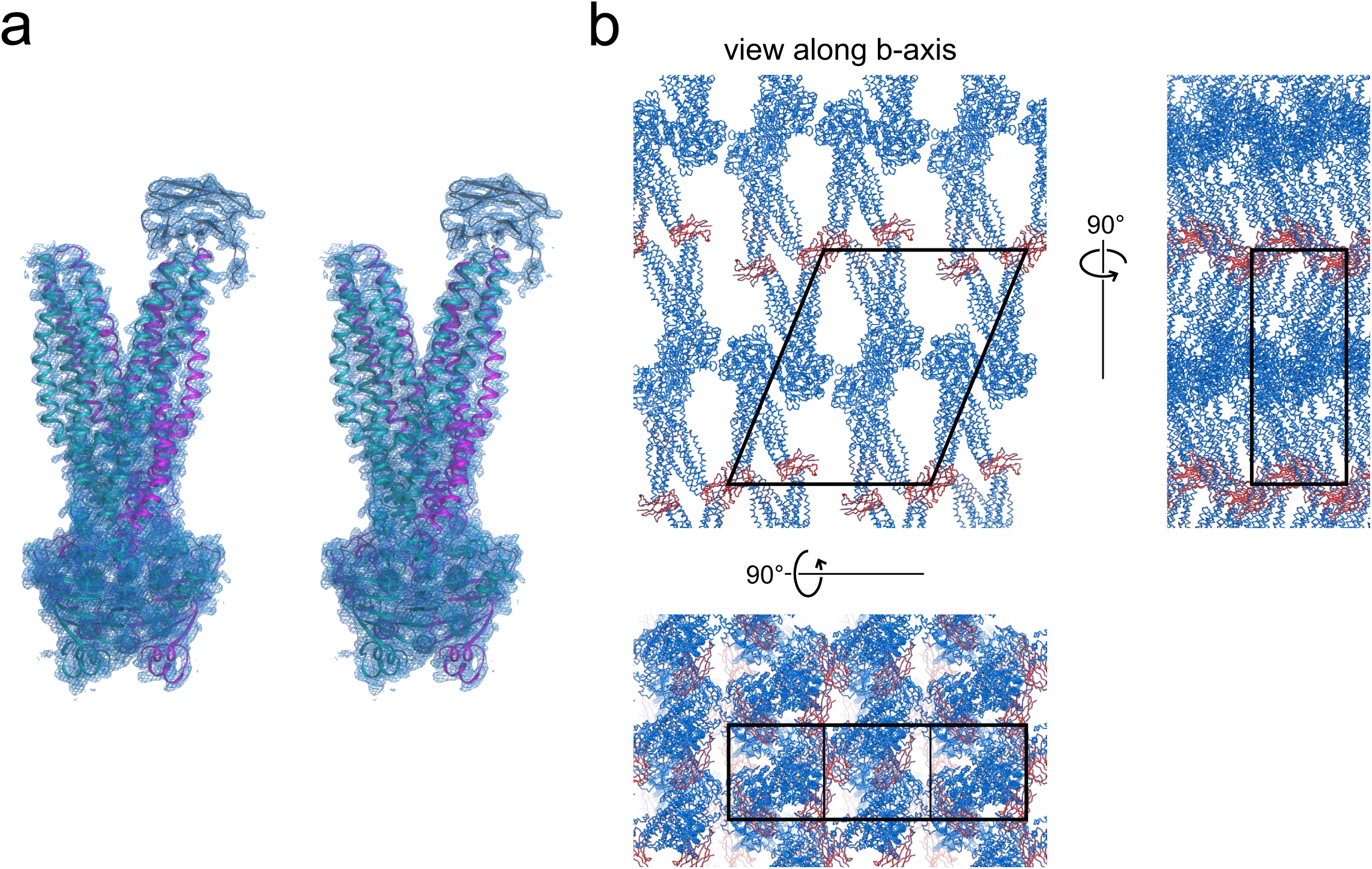
Electron density and crystal packing. (**a**) Stereo view of TM287/288–Sb_TM#35 complex structure. Electron density maps are shown as blue mesh contoured at 1.2 σ. (**b**) Sybody Sb_TM#35 (red) is involved in crystal packing.

**Figure S2.**
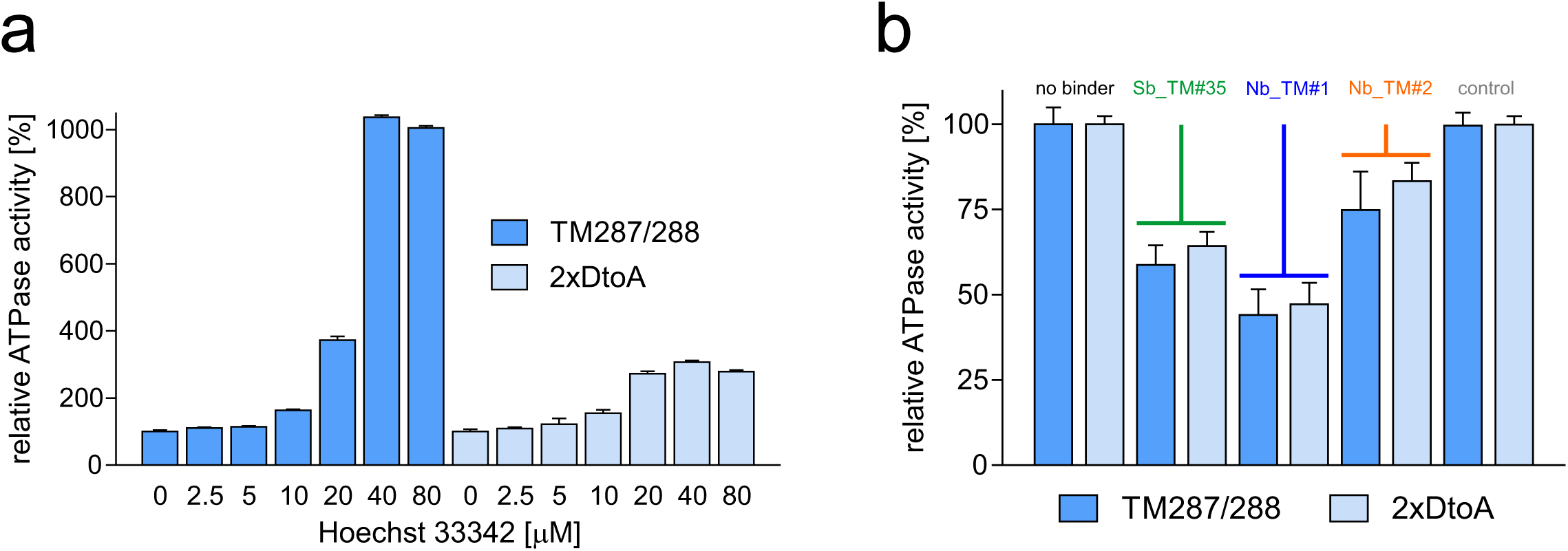
Characterization of TM287/288 reconstituted in nanodiscs. TM287/288 wildtype and 2xDtoA mutant were reconstituted into nanodiscs as described in Materials and Methods. (**a**) ATPase activity in the absence (basal activity) or presence of increasing Hoechst 33342 concentrations. Data were normalized to the basal activity of wildtype transporter or 2xDtoA mutant, respectively. (**b**) Inhibition of Hoechst 33342 stimulated ATPase activities (50 μM) in the presence of the binders Sb_TM#35, Nb_TM#1 or Nb_TM#2 (10 μM). A non-randomized sybody (10 μM) served as control. Data were normalized to the ATPase activity measured in the absence of binders. The error bars are standard deviations of technical triplicates.

**Figure S3.**
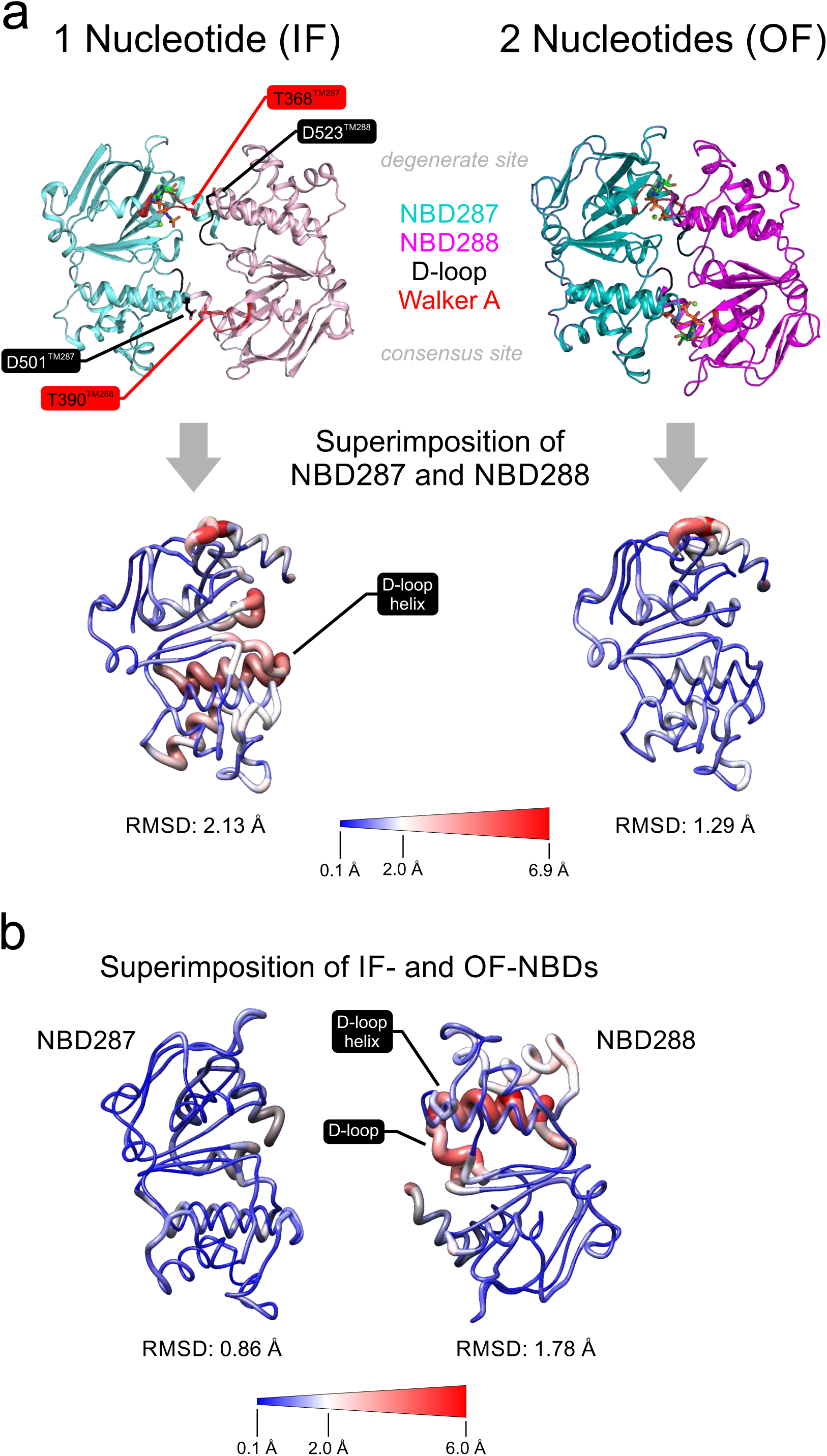
Superimposition of NBDs. Superimpositions are shown as sausage representations in which greater thickness and red coloring correlate with larger structural deviations. (**a**) Superimposition of NBD287 with NBD288 of inward-facing (IF) TM287/288 with bound AMP-PNP-Mg (left, PDB:4Q4H) or outward-facing (OF) with two bound ATPγS-Mg (right). Strong asymmetries found at the D-loop and D-loop helix of the IF structure are resolved in the closed NBD dimer of the OF structure. (**b**) Structural changes within the same NBDs during IF-OF conversion. While NBD287 moves as a rigid body, large rearrangements occur at the D-loop and D-loop helix of NBD288.

**Figure S4.**
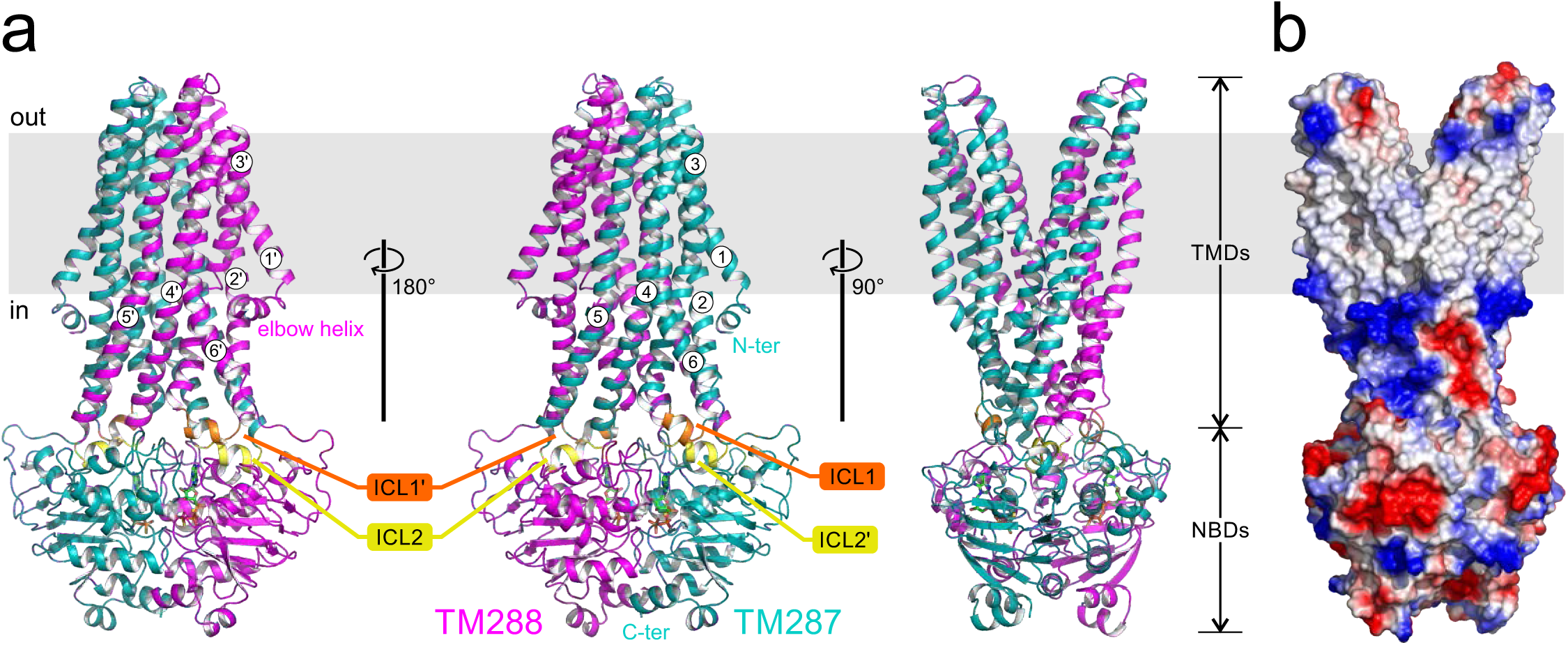
Side view of outward-facing TM287/288. (**a**) Cartoon representation of TM287/288 is colored in teal (TM287) and magenta (TM288) seen from different angles. The grey rectangle shows the membrane region. The twelve transmembrane helices are numbered (1-6 for TM287 and 1’-6’ for TM288). Intracellular loop 1 (ICL1) of TM287/288 features a kink that is not seen in ICL1’ of TM288 and is not common in ABC exporter structures. (**b**) Surface representation of outward-facing TM287/288 (positive and negative charges in blue and red, respectively). Hydrophobic and amphiphilic substrates could exit the binding cavity while remaining partially embedded in the outer leaflet of the lipid bilayer.

**Figure S5.**
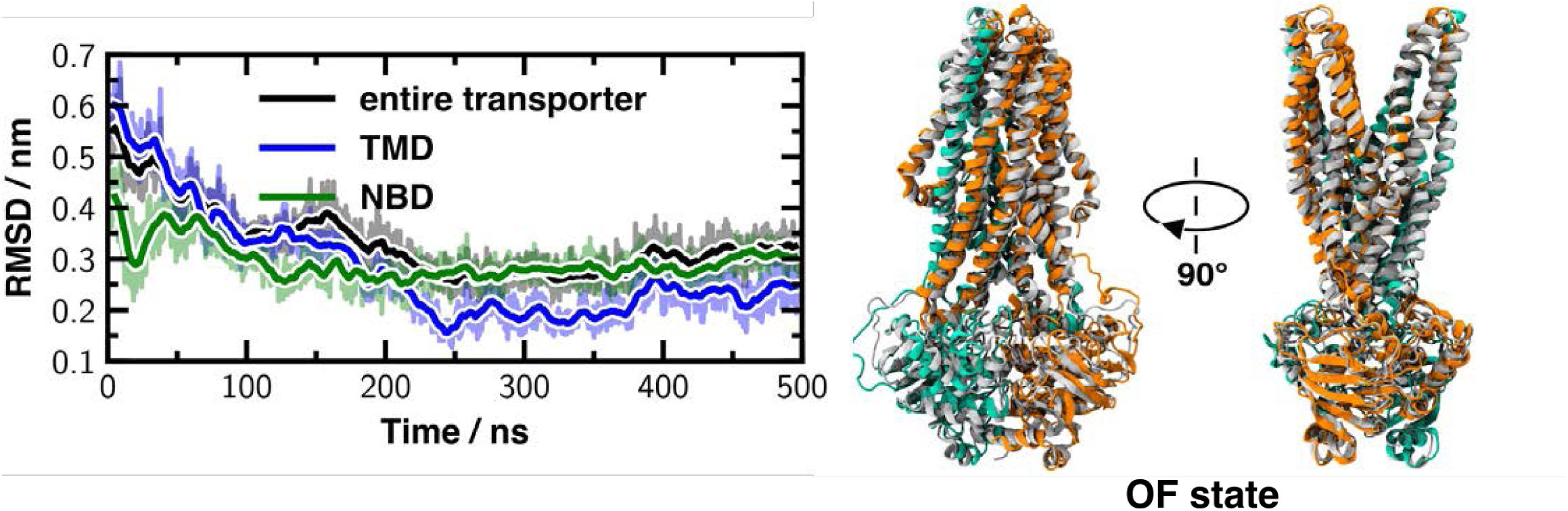
Comparison of outward-facing TM287/288 predicted by MD simulation to the crystal structure. Left panel: RMSD between C_α_-atoms of the OF crystal structure and the coordinates of the MD simulation during a representative IF-OF transition. Right panel: Superimposition of the predicted structure after 500 ns of MD simulation (TM287: orange, TM288: cyan) with the crystal structure of outward-facing TM287/288 (grey).

**Figure S6.**
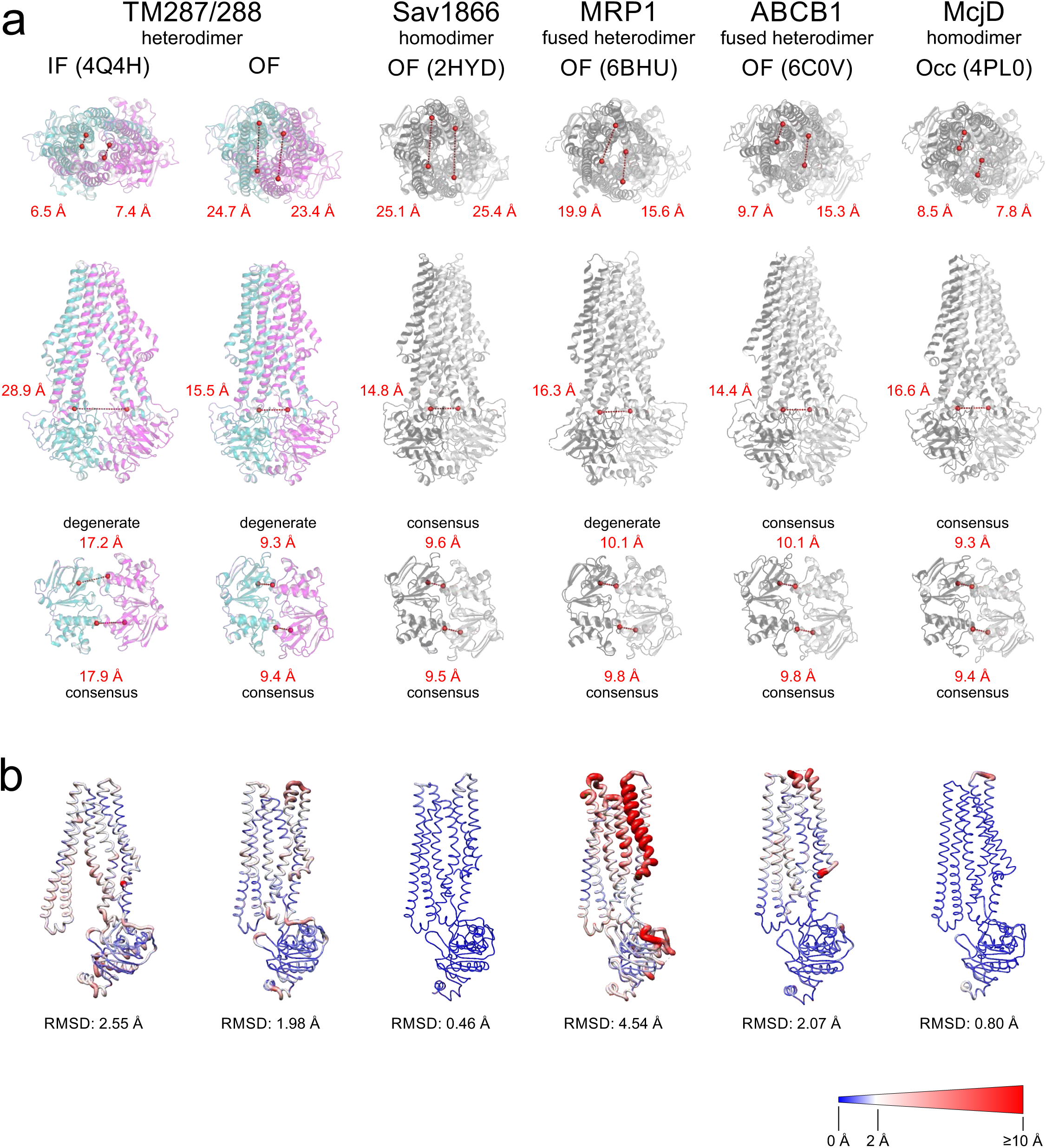
Asymmetries of outward-facing and outward-occluded ABC exporters. High resolution structures of type I ABC exporters were analyzed with regard to asymmetries among the respective half-transporters. (**a**) Distances (C_α_-C_α_) were measured in the extracellular gate (top panel, residues corresponding to D41^TM287^-E268^TM287^ and D65^TM288^-T292^TM288^, respectively), at the tetrahelix bundle (middle panel, residues corresponding to G201^TM287^-G225^TM288^), and the NBDs (bottom panel, distance between Walker A lysine and ABC signature serine in degenerate and consensus site, respectively). (**b**) Superimposition of the half-transporters shown in sausage representation. Greater thickness and red coloring correlate with larger structural deviations between equivalent positions.

**Figure S7.**
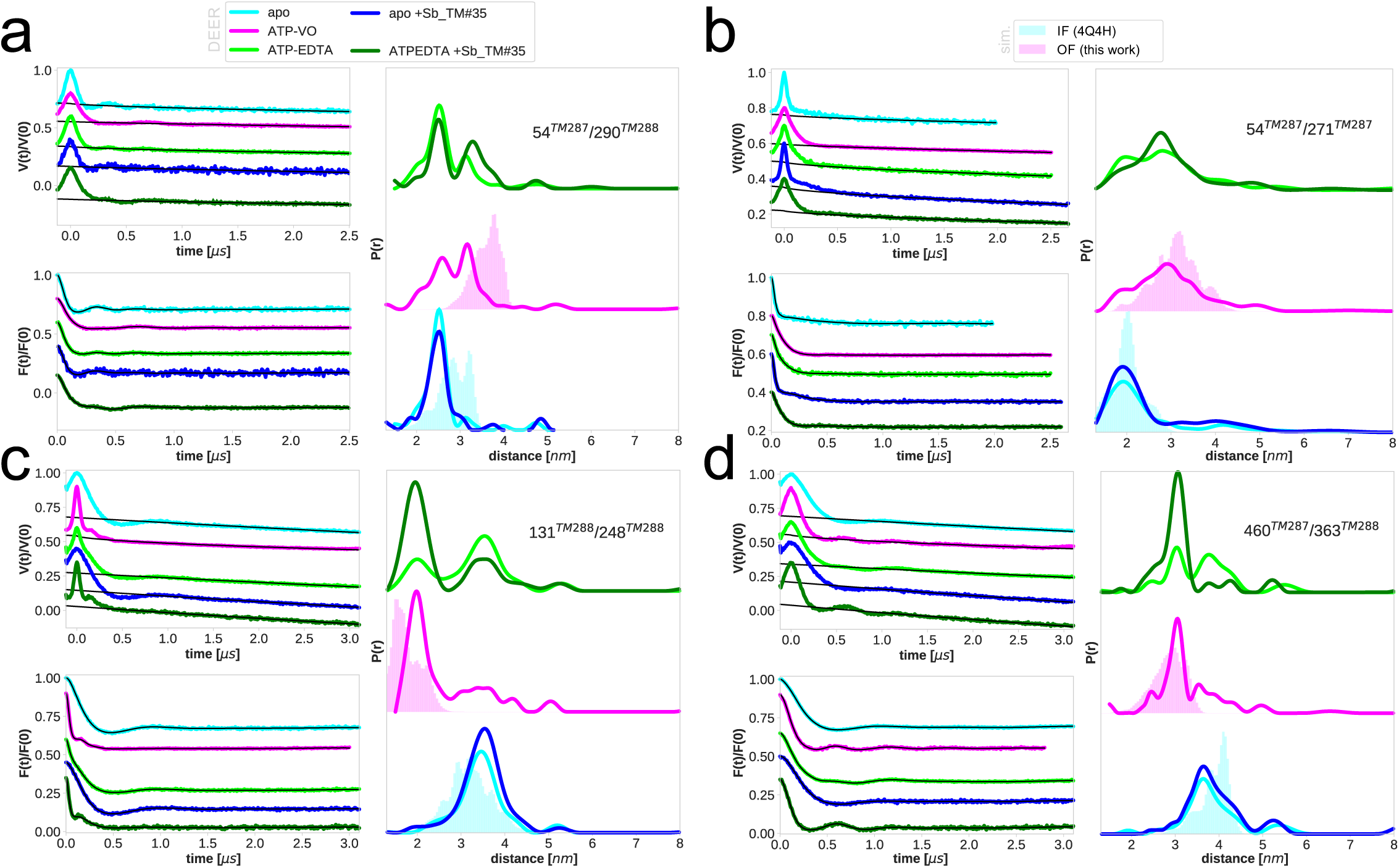
Primary DEER traces of spin-labeled pairs in TM287/288 measured in the presence or absence of unlabeled Sb_TM#35. Each panel shows Q-band DEER traces [V(t)/V(0)] with the fitted background, background-corrected [F(t)/F(0)] traces with fitted distribution function and the corresponding area-normalized distance distributions calculated using DeerAnalysis2015. Measurements were conducted in the absence of nucleotide (apo) or in the presence of ATP and vanadate (ATP-VO) or ATP-EDTA. Distance distributions were simulated using the program MMM2015 based on the coordinates of the IF and OF structures of TM287/288. (**a** and **b**) Extracellular pairs 54^TM287^/290^TM288^ and 54^TM287^/271^TM287^. (**c**) Intracellular pair 131^TM288^/248^TM288^. (**d**) NBD pair 460^TM287^/363^TM288^.

**Figure S8.**
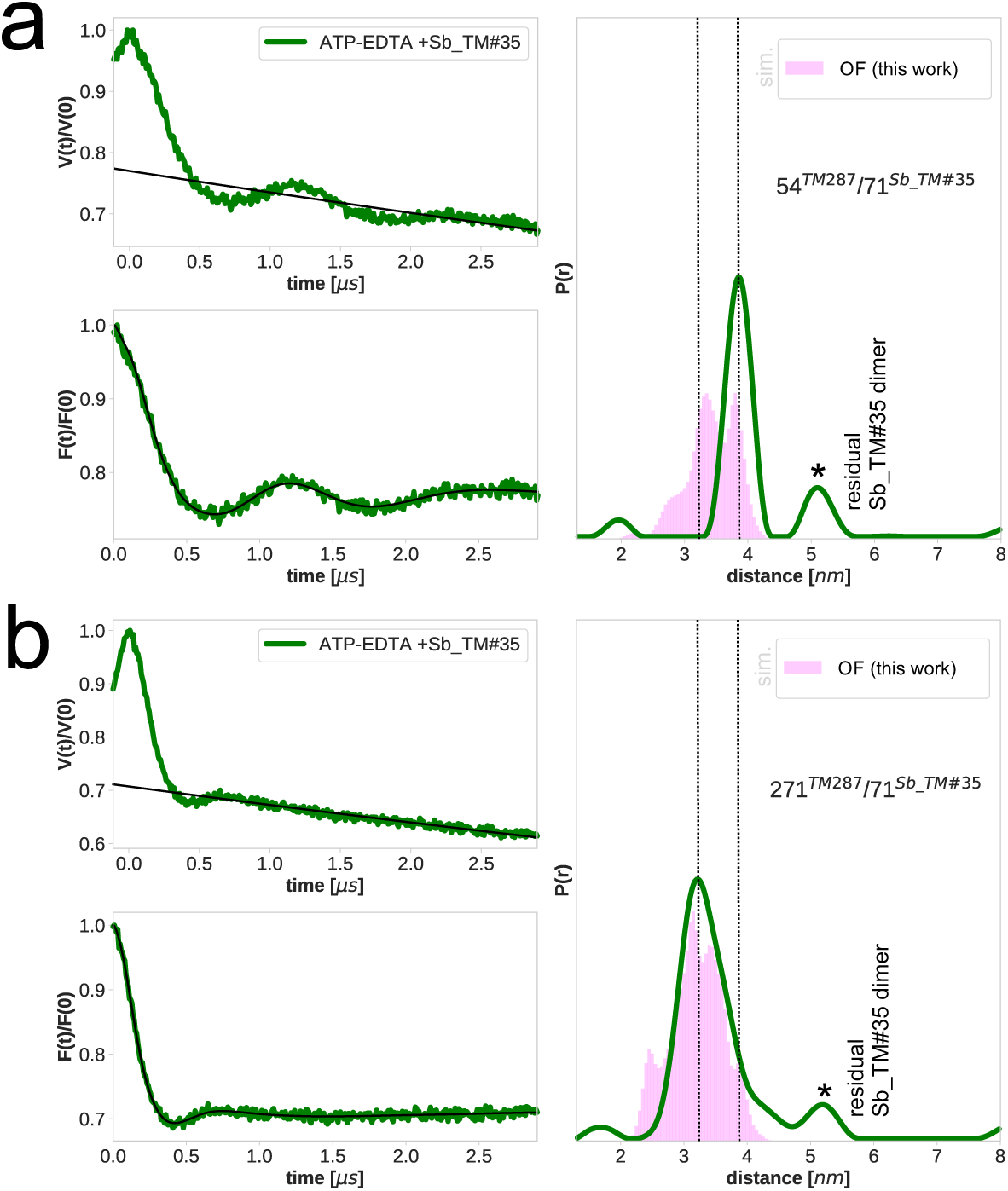
Primary DEER traces of spin-labeled pairs in TM287/288 and Sb_TM#35. Each panel shows Q-band DEER traces [V(t)/V(0)] with the fitted background, background-corrected [F(t)/F(0)] traces with fitted distribution function and the corresponding area-normalized distance distributions calculated using DeerAnalysis2015. Measurements were conducted in the presence of ATP-EDTA. Distance distributions were simulated using the program MMM2015 based on the coordinates of the OF structure of TM287/288 in complex with Sb_TM#35. (**a**) DEER analysis of 71^Sb_TM#35^ and 54^TM287^ positioned on the same wing where Sb_TM#35 binds. (**b**) DEER analysis of 71^Sb_TM#35^ and 271^TM287^ positioned on the opposite wing. Asterisks denote the residual 5.2 nm peak confirmed to arise from sybody dimers in solution, which can be detected as a unique peak in the absence of ATP, as well as in the absence of transporter (data not shown).

**Figure S9.**
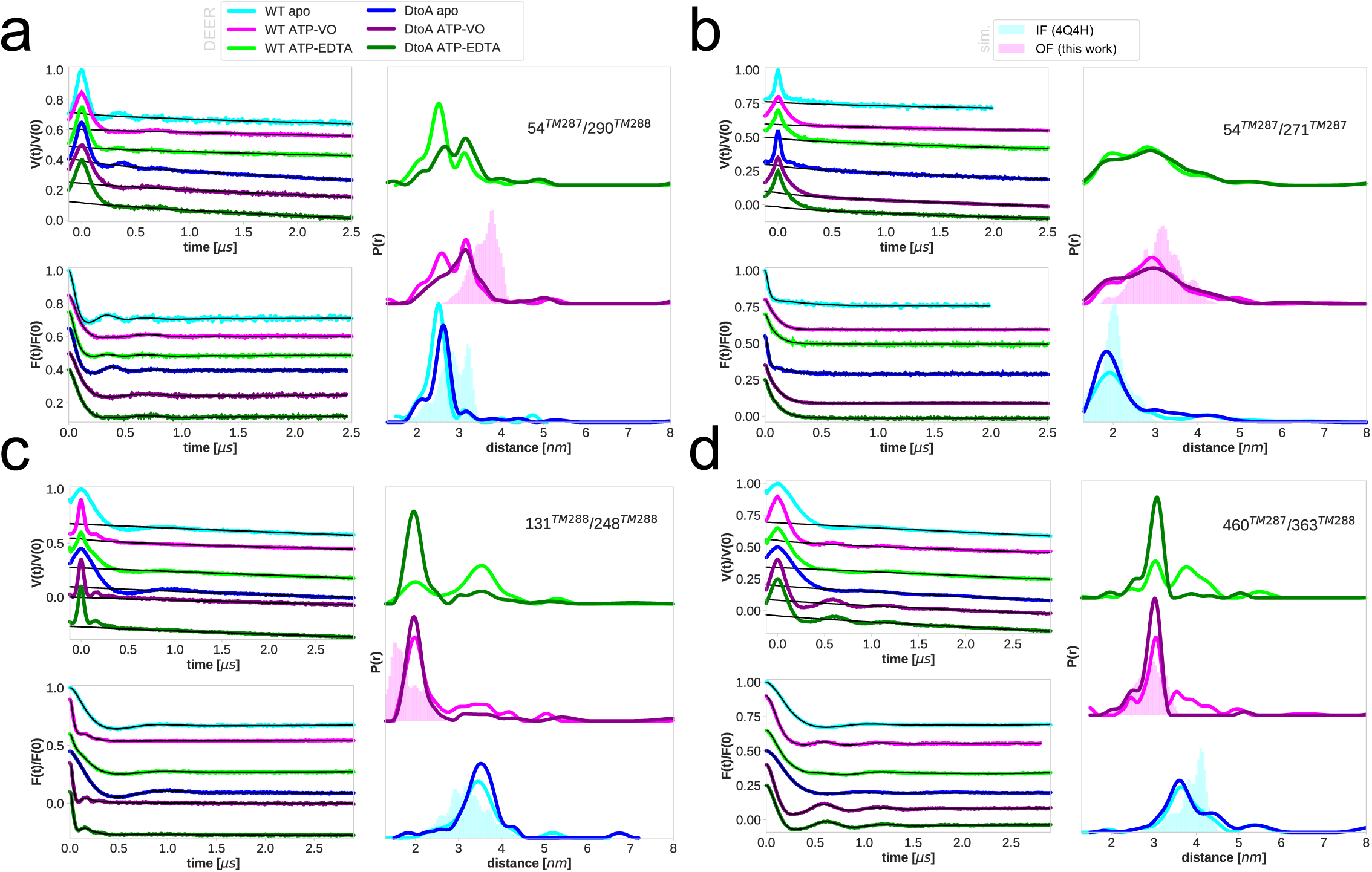
Primary DEER traces of spin-labeled pairs of wildtype TM287/288 and TM287/288(2xDtoA). Each panel shows Q-band DEER traces [V(t)/V(0)] with the fitted background, background-corrected [F(t)/F(0)] traces with fitted distribution function and the corresponding area-normalized distance distributions calculated using DeerAnalysis2015. Measurements were conducted in the absence of nucleotide (apo) or in the presence of ATP and vanadate (ATP-VO) or ATP-EDTA. Distance distributions were simulated using the program MMM2015 based on the coordinates of the IF and OF structures of TM287/288. (**a** and **b**) Extracellular pairs 54^TM287^/290^TM288^ and 54^TM287^/271^TM287^. (**c**) Intracellular pair 131^TM288^/248^TM288^. (**d**) NBD pair 460^TM287^/363^TM288^.

**Figure S10.**
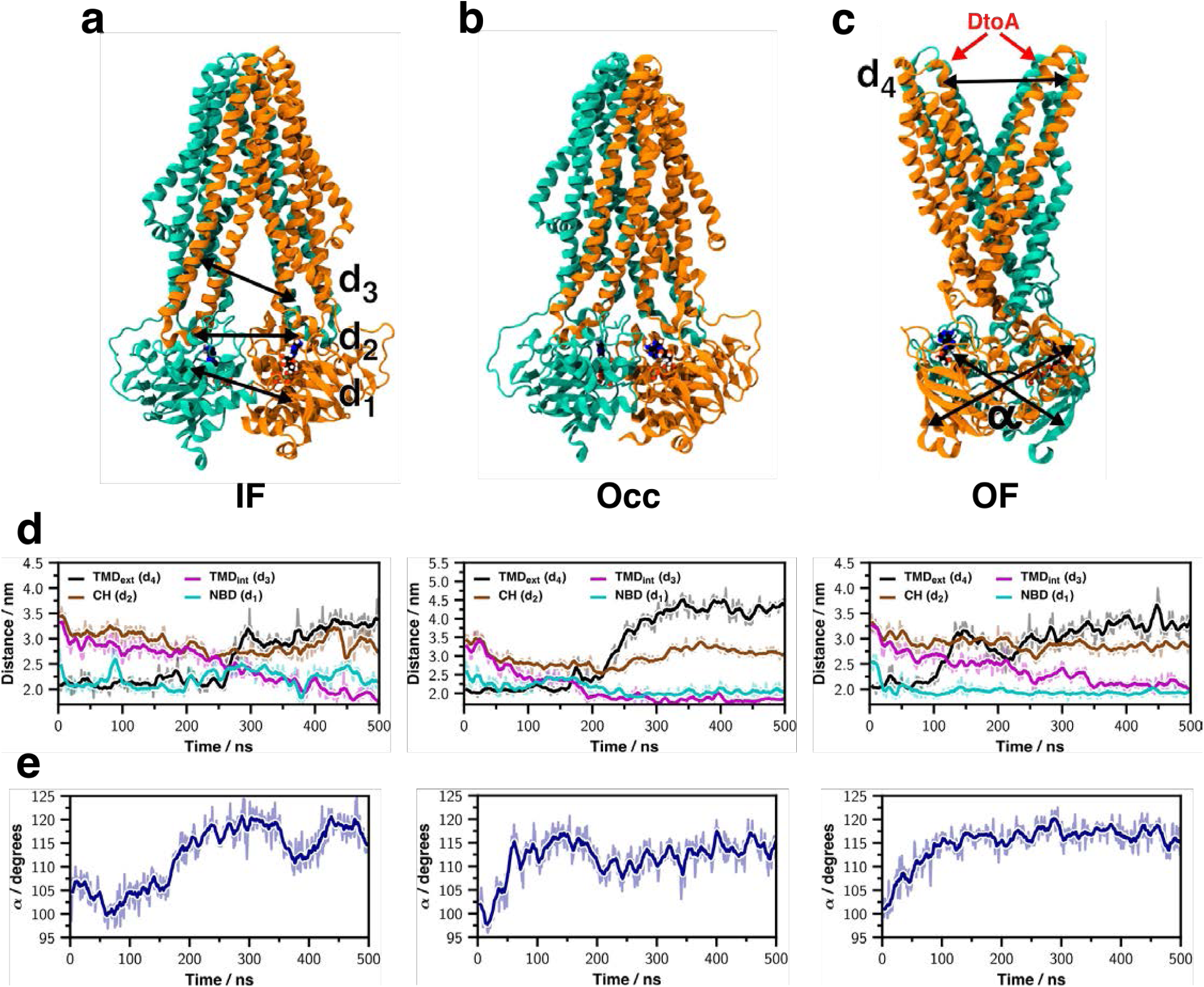
IF-OF transition of TM287/288(2xDtoA) captured by MD simulations. (**a**-**c**) IF, Occ, and OF conformations sampled during MD simulation. C_α_-C_α_ distances are measures for NBD dimerization (d_1_: D460^TM287^-S363^TM288^), coupling helix motion (d_2_: F127^TM288^-T227^TM288^), cytoplasmic gate closure (d_3_: T131^TM288^-S248^TM288^) and extracellular gate opening (d_4_: S50^TM287^-S271^TM287^). The arrows in (**c**) indicate the positions of mutated aspartates in the extracellular gate. (**d**) Distance time traces during three successful IF-to-OF conformational transitions. (**e**) Corresponding time traces of NBD-NBD twist angle, defined as the angle between the 2 vectors connecting C_α_-atoms of L554^TM287^-I452^TM287^ and L576^TM288^-I474^TM288^, respectively.

**Figure S11.**
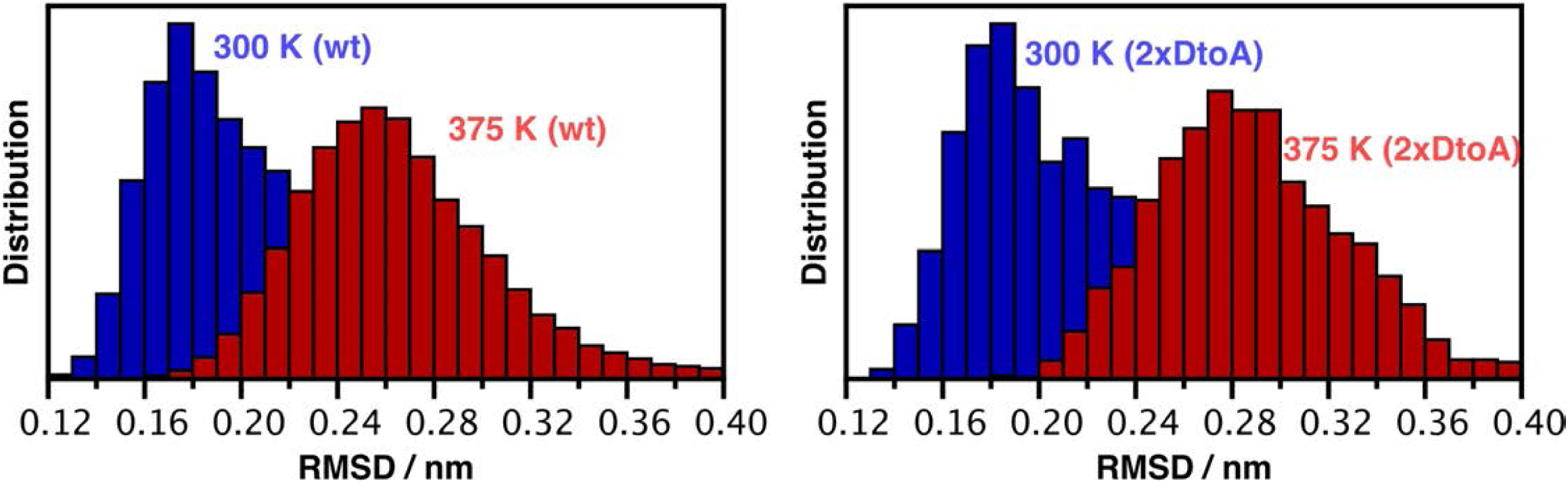
Stability of OF crystal structure during MD simulations. C_α_-RMSD distribution from MD simulations of wildtype (right panel) and 2xDtoA mutant (right panel). Ten MD simulations (of 400 ns each) were carried out for both wildtype and 2xDtoA mutant at 300 K (blue) and at 375 K (red), respectively (i.e., 40 simulations in total).

**Figure S12.**
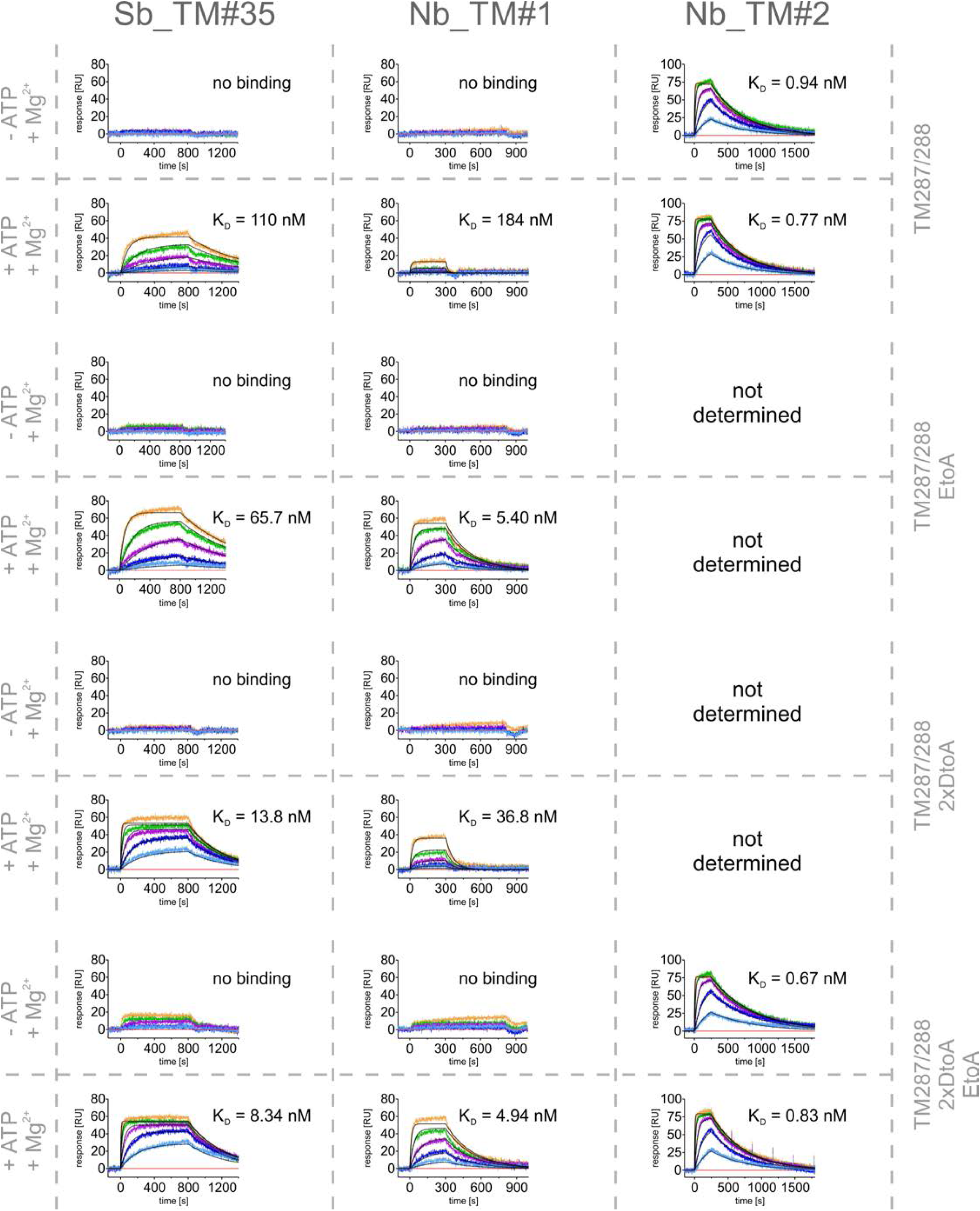
SPR analysis of single domain antibodies. TM287/288 and the respective mutants were immobilized on the SPR sensor chip and Sb_TM#35, Nb_TM#1 and Nb_TM#2 served as analytes. Binding experiments were conducted in an Mg^2+^-containing buffer in the presence or absence of ATP. Injected concentrations: Sb_TM#35: 0, 9, 27, 81, 243, 729 nM; Nb_TM#1: 0, 1, 3, 9, 27, 81 nM; Nb_TM#2: 0, 0.9, 2.7, 8.1, 24.3, 72.9 nM. Kinetic analyses are shown in Table S2.

**Figure S13.**
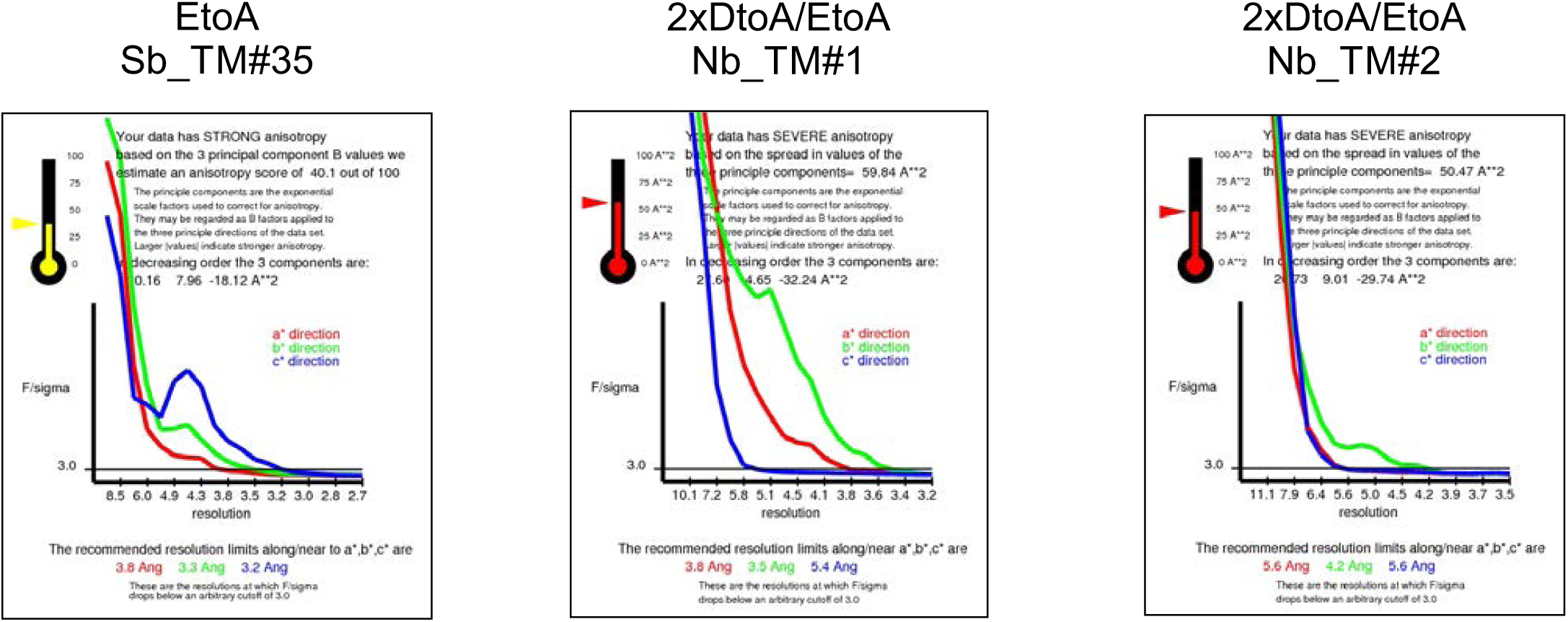
Crystallographic diffraction data truncation using the diffraction anisotropy server. Truncations were carried out using standard setting (F/sigma ≥ 3.0).

## TABLES

**Table S1.** Data collection and refinement statistics. Values in parentheses are for highest-resolution shell. #All datasets have been anisotropy corrected subsequently.

|  | EtoA<br>Sb_TM#35 | 2xDtoA/EtoA<br>Nb_TM#1 | 2xDtoA/EtoA<br>Nb_TM#2 |
| --- | --- | --- | --- |
| **Data collection<sup>#</sup>** |  |  |  |
| Space group | P2<sub>1</sub> | P2<sub>1</sub> | P1 |
| Cell dimensions |  |  |  |
| *a*, *b*, *c* (Å) | 165.35, 77.11, 207.02 | 90.75, 199.26, 113.16 | 88.60, 113.07, 126.89 |
| $\alpha$, $\beta$, $\gamma$ (°) | 90.00, 112.49, 90.00 | 90.00, 91.26, 90.00 | 83.18, 73.00, 67.37 |
| Resolution (Å) | 50-3.2 (3.28-3.20)<sup>*</sup> | 50-3.5 (3.59-3.50) | 50-4.2 (4.31-4.20) |
| *R*<sub>meas</sub> | 0.34 (3.91) | 0.09 (4.23) | 0.11 (3.11) |
| *I* / σ*I* | 7.28 (0.75) | 9.77 (0.63) | 8.89 (0.52) |
| Completeness (%) | 99.8 (99.7) | 98.8 (99.4) | 98.8 (98.7) |
| Redundancy | 13.6 (14.0) | 7.2 (7.4) | 3.5 (3.5) |
| CC<sub>1/2</sub> (%) | 99.8 (50.4) | 100 (40.2) | 100 (43.0) |
| **Refinement** |  |  |  |
| Resolution (Å) | 49.14-3.23 | 34.72-3.48 | 32.65-4.21 |
| No. reflections | 63'777 | 29'833 | 17'961 |
| *R*<sub>work</sub> / *R*<sub>free</sub> | 25.2/27.0 | 26.6/30.0 | 29.9/32.0 |
| No. atoms |  |  |  |
| Protein | 19'907 | 19'788 | 19'970 |
| Ligand/ion | 132 | 128 | 128 |
| Water | 0 | 0 | 0 |
| *B*-factors |  |  |  |
| Protein | 73.6 | 192.8 | 240.5 |
| Ligand/ion | 47.8 | 132.9 | 234.2 |
| R.m.s. deviations |  |  |  |
| Bond lengths (Å) | 0.008 | 0.008 | 0.008 |
| Bond angles (°) | 0.93 | 0.94 | 0.92 |

<sup>\*</sup>Values in parentheses are for highest-resolution shell.
<sup>#</sup>All datasets have been anisotropy corrected subsequently.

**Table S2.** Kinetic parameters derived from SPR measurements. x, no SPR signal; n.d., not determined

|  |  | Sb_TM#35 |  | Nb_TM#1 |  | Nb_TM#2 |  |
| --- | --- | --- | --- | --- | --- | --- | --- |
|  |  | - ATP<br>+ Mg<sup>2+</sup> | + ATP<br>+ Mg<sup>2+</sup> | - ATP<br>+ Mg<sup>2+</sup> | + ATP<br>+ Mg<sup>2+</sup> | - ATP<br>+ Mg<sup>2+</sup> | + ATP<br>+ Mg<sup>2+</sup> |
| **TM287/288** | $k_{on}$ (M<sup>-1</sup>s<sup>-1</sup>) | x | 1.43*10<sup>4</sup> | x | 3.41*10<sup>5</sup> | 2.18*10<sup>6</sup> | 2.77*10<sup>6</sup> |
| | $k_{off}$ (s<sup>-1</sup>) | x | 1.57*10<sup>-3</sup> | x | 6.28*10<sup>-2</sup> | 2.06*10<sup>-3</sup> | 2.12*10<sup>-3</sup> |
| | $K_D$ (M) | x | 1.10*10<sup>-7</sup> | x | 1.84*10<sup>-7</sup> | 9.43*10<sup>-10</sup> | 7.65*10<sup>-10</sup> |
| **TM287/288**<br>**EtoQ** | $k_{on}$ (M<sup>-1</sup>s<sup>-1</sup>) | x | 2.06*10<sup>4</sup> | x | 1.01*10<sup>6</sup> | 2.44*10<sup>6</sup> | 2.36*10<sup>6</sup> |
| | $k_{off}$ (s<sup>-1</sup>) | x | 1.35*10<sup>-3</sup> | x | 5.04*10<sup>-3</sup> | 1.96*10<sup>-3</sup> | 2.19*10<sup>-3</sup> |
| | $K_D$ (M) | x | 6.56*10<sup>-8</sup> | x | 4.98*10<sup>-9</sup> | 8.03*10<sup>-10</sup> | 9.28*10<sup>-10</sup> |
| **TM287/288**<br>**EtoA** | $k_{on}$ (M<sup>-1</sup>s<sup>-1</sup>) | x | 1.92*10<sup>4</sup> | x | 9.01*10<sup>5</sup> | n.d. | n.d. |
| | $k_{off}$ (s<sup>-1</sup>) | x | 1.26*10<sup>-3</sup> | x | 4.86*10<sup>-3</sup> | n.d. | n.d. |
| | $K_D$ (M) | x | 6.57*10<sup>-8</sup> | x | 5.40*10<sup>-9</sup> | n.d. | n.d. |
| **TM287/288**<br>**2xDtoA** | $k_{on}$ (M<sup>-1</sup>s<sup>-1</sup>) | x | 1.84*10<sup>5</sup> | x | 4.27*10<sup>5</sup> | n.d. | n.d. |
| | $k_{off}$ (s<sup>-1</sup>) | x | 2.55*10<sup>-3</sup> | x | 1.57*10<sup>-2</sup> | n.d. | n.d. |
| | $K_D$ (M) | x | 1.38*10<sup>-8</sup> | x | 3.68*10<sup>-8</sup> | n.d. | n.d. |
| **TM287/288**<br>**EtoA**<br>**2xDtoA** | $k_{on}$ (M<sup>-1</sup>s<sup>-1</sup>) | x | 2.82*10<sup>5</sup> | x | 8.92*10<sup>5</sup> | 2.40*10<sup>6</sup> | 2.44*10<sup>6</sup> |
| | $k_{off}$ (s<sup>-1</sup>) | x | 2.36*10<sup>-3</sup> | x | 4.40*10<sup>-3</sup> | 1.61*10<sup>-3</sup> | 2.03*10<sup>-3</sup> |
| | $K_D$ (M) | x | 8.34*10<sup>-9</sup> | x | 4.94*10<sup>-9</sup> | 6.73*10<sup>-10</sup> | 8.29*10<sup>-10</sup> |

**Table S3.** ATPase activities of spin-labeled TM287/288 determined using 500 µM ATP at 25°C.

| | ATPase activity<br>(nmol Pi / min / mg protein) | Inhibited ATPase activity<br>+ 1.2 μM Sb_TM#35<br>(nmol Pi / min / mg protein) |
| --- | --- | --- |
| wildtype | 97.6 ± 6.7 | 21.7 ± 1.3 |
| 54<sup>TM287</sup>/290<sup>TM288</sup> | 59.4 ± 3.7 | 22.1 ± 2.0 |
| 54<sup>TM287</sup>/271<sup>TM287</sup> | 44.5 ± 3.6 | 14.1 ± 1.7 |
| 54<sup>TM287</sup> | 78.3 ± 4.4 | 23.7 ± 2.7 |
| 271<sup>TM287</sup> | 48.2 ± 3.0 | 21.3 ± 2.1 |

